# Cold-induced expression of a truncated Adenylyl Cyclase 3 acts as rheostat to brown fat function

**DOI:** 10.1101/2022.08.01.502156

**Authors:** Sajjad Khani, Hande Topel, Ajeetha Josephrajan, Bjørk Ditlev Marcher Larsen, Ana Rita Albuquerque de Almeida Tavanez, Michael James Gaudry, Philipp Leyendecker, Natasa Stanic, Isabella Gaziano, Nils Rouven Hansmeier, Elena Schmidt, Paul Klemm, Lara-Marie Vagliano, Christoph Andreas Engelhard, Søren Nielsen, Naja Zenius Jespersen, Rizwan Rehimi, Sabrina Gohlke, Peter Frommolt, Thorsten Gnad, Alvaro Rada-Iglesias, Marta Pradas-Juni, Tim Julius Schulz, Frank Thomas Wunderlich, Alexander Pfeifer, Martin Jastroch, Dagmar Wachten, Jan-Wilhelm Kornfeld

**Affiliations:** Institute for Genetics, University of Cologne, Zülpicher Str. 47a, 50674 Cologne, Germany; Max-Planck Institute for Metabolism Research, Gleueler Strasse 50, 50931 Cologne, Germany; Department for Biochemistry and Molecular Biology, University of Southern Denmark, Campusvej 55, 5230 Odense, Denmark; Department of Molecular Biosciences, The Wenner-Gren Institute, Stockholm University, SE-106 91, Stockholm, Sweden; Institute of Innate Immunity, Medical Faculty, University of Bonn, Venusberg-Campus 1, 53127 Bonn, Germany; Centre for Physical Activity Research, Department of Infectious Diseases, Rigshospitalet, Faculty of Health Sciences, University of Copenhagen, Blegdamsvej 9, 2100 Copenhagen, Denmark; Center for Molecular Medicine Cologne (CMMC), University of Cologne, Robert-Koch-Str. 2150931 Cologne, Germany; Department of Adipocyte Development and Nutrition, German Institute of Human Nutrition Potsdam-Rehbrücke, Arthur-Scheunert-Allee 114-116, 14558 Nuthetal, Germany; Institute of Human Genetics, University Medical Center Hamburg-Eppendorf, Martinistr. 52, 20246 Hamburg, Germany; Institute of Pharmacology and Toxicology, University Hospital, University of Bonn, Venusberg-Campus 1, 43127, Bonn, Germany; Institute of Biomedicine and Biotechnology of Cantabria (IBBTEC), CSIC/University of Cantabria, C/ Albert Einstein 22, 39011 Santander; Novo Nordisk Foundation Center for Basic Metabolic Research (CBMR), Blegdamsvej 3a, 2200 Copenhagen, Denmark

**Author notes:** These authors contributed equally to this work (Sajjad Khani, Hande Topel, Dagmar Wachten, Jan-Wilhelm Kornfeld).

## Abstract

Promoting brown adipose tissue (BAT) activity has been recognized as innovative therapeutic approach to improve obesity and metabolic disease. Whilst the molecular circuitry underlying thermogenic activation of BAT is well understood, the processes underlying rheostatic regulation of BAT to maintain homeostasis and avoid excessive energy dissipation remain ill-defined. Increasing cyclic AMP (cAMP) biosynthesis is key for BAT activation. Here, we demonstrate that ADCY3, an adenylyl cyclase whose expression is induced during cold exposure and regulates cAMP homeostasis in thermogenic fat, is dispensable for BAT function in lean mice, but becomes critical during obesity. Furthermore, by combining RNA-seq with epigenomic H3K4me3 profiling, we detected a novel, cold-inducible promoter that generates a 5’ truncated *Adcy3-at* mRNA isoform, *Adcy3-at*. Mice lacking only *Adcy3-at*, but not *full-length Adcy3*, displayed increased energy expenditure already under lean conditions and were protected against obesity and ensuing metabolic imbalances. Subcellularly, translated ADCY3-AT proteins are retained in the endoplasmic reticulum (ER), did not translocate to the cell membrane, and lacked enzymatic activity. By interacting with ADCY3, ADCY3-AT retained ADCY3 in the ER and, thereby, reduced the plasma membrane pool of ADCYs available for G-protein mediated cAMP synthesis. Thereby, ADCY3-AT acts as a signaling rheostat in BAT, limiting adverse consequences of uncurbed cAMP activity after long-term BAT activation. *Adcy3-at* induction was driven by a cold-induced, truncated isoform of the transcriptional cofactor PPARGC1A (PPARG Coactivator 1 Alpha, PPARGC1A-AT). Expression of *Ppargc1a-at* and *Adcy3-at* are evolutionary conserved, indicating that transcriptional rewiring by commissioning of alternative promoters is key for thermogenic fat function.

## Introduction

Most mammals harbor two morphologically and functionally distinct types of adipocytes (fat cells): White adipocytes consist of a single lipid droplet^1^ and predominantly regulate energy storage, whereas brown adipocytes are multilocular and possess the ability to convert (diet-derived) macronutrients like carbohydrates and lipids into heat in a unique molecular process termed Non-Shivering Thermogenesis (NST)^2^. Interest in brown adipose tissue (BAT) arose after the seminal discovery that *(i)* BAT not only exists in small animals like rodents and in human infants, but that adult humans can possess active brown fat ^3–5^ and *(ii)* cold ambient temperature exposure or stimulation of beta-adrenergic G-Protein receptors (GPCRs) activates BAT, which positively correlates with energy expenditure^6^ and inversely correlates with body-mass indices^7, 8^.

GPCRs regulate a plethora of biological processes and are the target of a major part of drugs in clinical use^9^. GPCRs play a central role in adipose tissue homeostasis, and adipocyte metabolism is regulated by GPCRs linked to different functional classes of heterotrimeric G proteins ^10^. The main physiological stimulus for NST in brown and thermogenically activated ‘brite’ (brown-in-white) fat is cold temperature, which activates the sympathetic nervous system (SNS) and leads to release of the neurotransmitter norepinephrine (NE) ^11^. NE activates beta-adrenergic GPCRs^12^, which then stimulate synthesis of the second-messenger 3’-5’-cyclic adenosine monophosphate (cAMP) by plasma membrane-bound adenylyl cyclases^13, 14^ and activates protein kinase A (PKA), thereby inducing lipolysis. The ensuing generation of free fatty acids (FFA) activates uncoupling protein 1 (UCP1), a mitochondrial protein whose activation executes NST in brown and brite fat^2^. In addition to these acute effects, chronic cAMP signaling is required for adipocyte precursor differentiation ^2^. Adenylyl cyclase 3 (ADCY3), which is expressed in somatic tissues like adipose tissue, kidney, pancreas and liver as well as in olfactory sensory and hypothalamic neurons^15^, received attention due to loss-of-function variants in human *ADCY3* that correlate with enhanced susceptibility to increased BMI^16^ and *ADCY3* loss-of-function mutations cause severe obesity in multiple human populations ^17–19^. Mouse studies confirmed that *Adcy3* silencing is sufficient to impair energy homeostasis ^20–24^, yet the precise role of ADCY3 in thermogenic fat has not been addressed yet.

Pre-mRNA splicing is a fundamental gene-regulatory mechanism that enables cells to increase proteome complexity from a finite amount of exonic information and is important for adipocyte function, cold tolerance, and metabolic health ^25–27^. For instance, PPARG coactivator 1 alpha (PPARGC1A), a key transcriptional regulator in adipose tissue, displays an alternative 5’ promoter in human muscle, generating a C-terminally truncated protein that elicits distinct transcriptional programs ^28, 29^. It remains elusive though if similar alternative promoter events contribute to metabolic regulation in thermogenic fat. *Here, we describe a novel molecular rheostat, ADCY3-AT, that is selectively induced and controls brown fat function upon cold and beta-adrenergic stimulation. N-terminally truncated ADYC3-AT proteins limit cAMP synthesis by controlling the subcellular localization of adenylyl cyclases and ultimately energy dissipation. Truncated ADCY3 is evolutionary conserved in brown adipocytes from rodents to humans and is part of a conserved transcriptional network, linking thermogenic activation to adaptive proteoform changes*.

## Results

### Transcriptional regulation of cAMP biosynthesis during cold adaptation of brown adipose tissue

To identify transcriptional processes controlling 3’,5’-cyclic adenosine monophosphate (cAMP) second messenger production in cold-activated brown (BAT) and epididymal white adipose tissue (eWAT), we exposed chow diet-fed, male C57BL/6N mice to 24 hours of 5°C cold exposure (CE) and analyzed transcriptome-wide gene expression changes using mRNA-sequencing (RNA-seq). We combined our analysis with public mRNA-seq data from 72 h cold-exposed, inguinal white adipose tissue (iWAT)^30^ and observed significantly (FDR ≤ 0.05) up-or downregulated genes, respectively: 1,086/1,204 genes in BAT, 804/448 genes in iWAT, and 584/71 in eWAT (Fig.1a-c, Fig.S1a-c) genes between cold and ambient room (22°C) temperatures (s. Supplemental table S1). KEGG gene ontology (GO) analysis revealed that peroxisome and PPAR signaling, as well as protein metabolism processes were induced, whereas HIF-1-dependent hypoxic signaling was repressed in cold-activated BAT (Fig.1d). Canonical thermogenic and brown-fat associated GO terms like oxidative phosphorylation, thermogenesis, tricarbon acid cycle, and carbon metabolism were upregulated in iWAT (Fig.1e), whereas few categories were altered in eWAT (Fig.1f), likely reflecting the limited capacity for thermogenesis in this depot. We next specifically investigated the expression of cAMP-degrading and synthesizing enzymes: Whilst expression of negative regulators of cyclic nucleotides, namely 3′,5′-cyclic nucleotide phosphodiesterases (PDEs)^31^, remained unchanged across all depots investigated (Fig.S1d-f), mRNA levels of adenylyl cyclase isoforms 3 and 4 (*Adcy3*, *Adcy4*) were specifically induced in cold-exposed BAT (Fig.1g) and only *Adcy3* expression was induced in iWAT (Fig.1h), whereas expression of *Adcy* isoforms remained unaltered in eWAT (Fig.1i). These results point towards a specific role for ADCY3 in thermogenic adipocytes and regulation of energy expenditure as proposed previously ^32, 33^.

**Figure 1:**
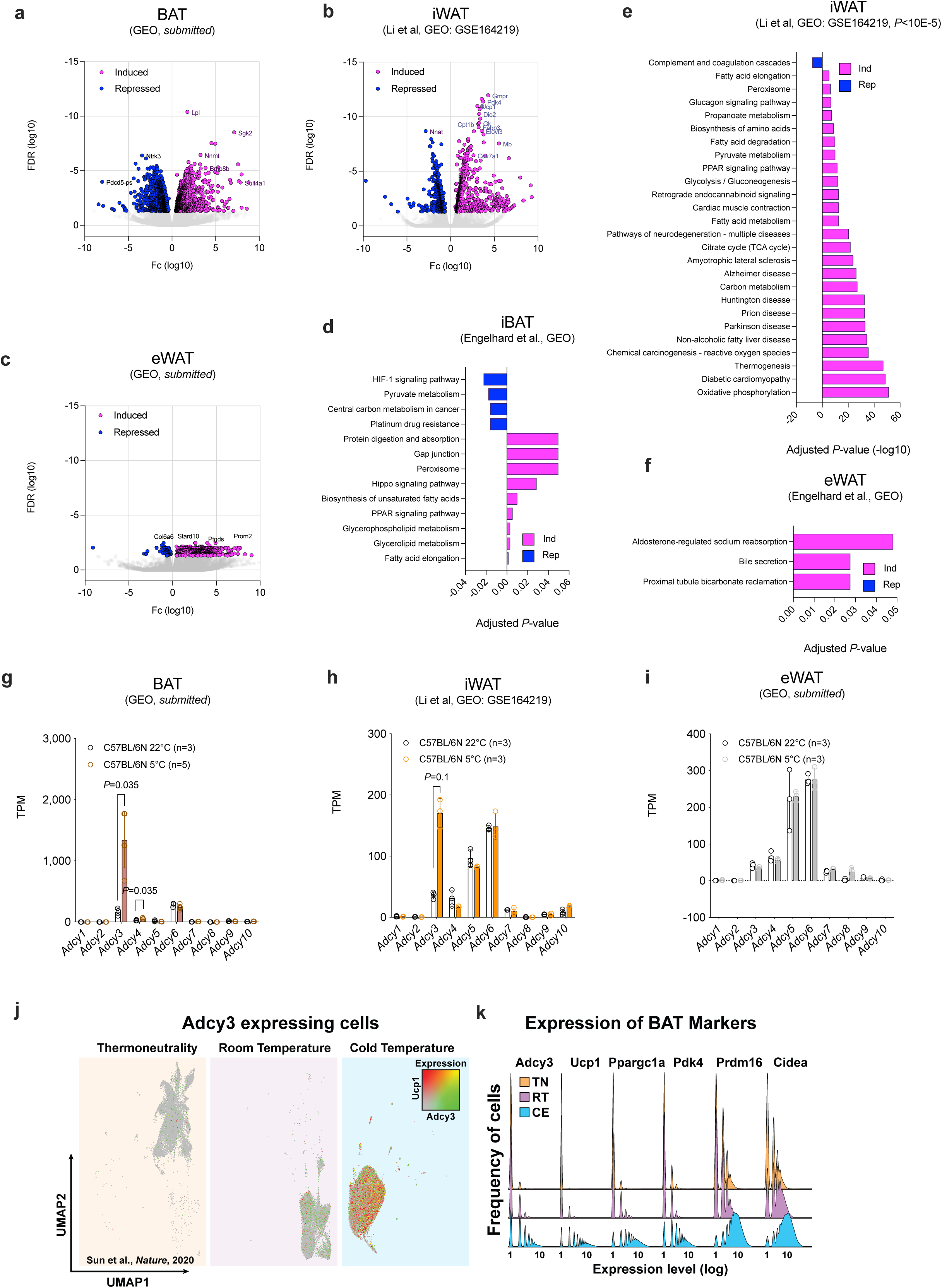
Transcriptional regulation of cAMP biosynthesis during cold adaptation of adipose tissue. (**a-c**) Volcano plot of all and significantly altered (cyan, magenta) genes in BAT (**a**, n=2,290; 1,086/1,204 up-/downregulated DEGs), iWAT (**b**, n=1,252; 804/448 up-/downregulated DEGs) and eWAT (**c**, n=655 up-/downregulated DEGs) in 20 weeks old C57BL/6N male mice housed at room temperature (22°C) or after 24 h (**a,c**) and 72 h (**b**) of 5°C cold exposure as determined by mRNA-seq. Statistically significant genes were defined as FDR ≤0.05 and log transcripts per million (TPM)>0. n=3-5 animals per temperature condition were analyzed. (**d-f**) KEGG pathway enrichments (Benjamini & Hochberg corrected *P*-value ≤ 0.05) for DEGs shown in (**a-c**). (**g-i**) mRNA expression of adenylyl cyclase isoforms abundance in BAT (**g**), iWAT (**h**) and eWAT (**i**) determined by mRNA-seq and depicted in TPM. Bar graphs represent mean ± SD with all data points plotted and unpaired, two-tailed, and non-parametric Mann-Whitney tests were performed to assess statistical significance. *P*-values are indicated within the panel. **(j-k**) Analysis of single-nucleus RNA-seq (snRNA-seq) data from *Adipoq*-tdTomato positive adipocyte nuclei housed at thermoneutrality (TN), room temperature (RT), and animals after cold exposure (CE) at 8°C for 4 days (**j**). The colors depict expression of *Adcy3* (green), *Ucp1* (red) or both (yellow). (**k**) Ridgeline plots depicting frequency (in nuclei) of expression of BAT identity genes at indicated temperatures.

Analysis of public single-nuclei RNA-Seq (snRNA-seq) datasets^34^ from purified brown adipocytes (BA) revealed that *Adcy3* is expressed lowly under basal (thermoneutral) conditions not exhibiting substantial NST. However, *Adcy3* expression is induced when lowering ambient temperatures from thermoneutrality (TN) to room temperature (RT) to cold exposure (CE) in brown adipocytes (BAs), paralleling increased expression changes of known BAT markers like uncoupling protein 1 (*Ucp1*, Fig.1j). Similar patterns were observed for known BAT markers^35^ like PPARGC1A, pyruvate dehydrogenase kinase 4 *(Pdk4)*, PR/SET domain 16 *(Prdm16),* and cell death inducing DFFA like effector A *(Cidea*, Fig.1k). Thus, cold activation induces *Adcy3* expression in cold-activated BA and inguinal white adipocytes (iWA).

### ADCY3 is required for cAMP biosynthesis in BAT and required for cold adaptation in obesity

To decipher the adipocyte-autonomous roles of ADCY3 *in vivo*, we next crossed mice harboring *LoxP*-flanked *Adcy3* alleles^24^ with animals expressing Adiponectin (*Adipoq*)-promoter driven cre recombinase^36^ (*Adipoq*-*cre*) to obtain pan-adipocyte-deficient *Adcy3* knockout mice (*Adcy3^LoxP/LoxP^, Adipoq-cre^+/cre^*; *Adcy3-AdcKO)* and cre-negative littermates *(Adcy3^LoxP/LoxP^, Adipoq-cre^+/+^*; *LoxP*, Fig.2a). RNA-seq demonstrated 60-70% reduction of *Adcy3* in *Adcy3-AdcKO* BAT with remaining *Adcy3* likely representing non-adipocyte cell populations in BAT^34^ (Fig.2b). Cyclic AMP levels in BAT of mice housed at room temperature were concomitantly reduced 50 %, pointing to ADCY3 constituting a pivotal cAMP regulator in thermogenic fat (Fig.2c). To probe for roles of adipocyte ADCY3 in energy metabolism, we analyzed metabolic parameters in lean, chow diet-fed *Adcy3-AdcKO* mice. Unexpectedly, we observed no differences in body weight (BW, Fig.2d), and intraperitoneal glucose (Fig.2e) and insulin (Fig.2f) tolerance tests revealed no differences in glucose metabolism between chow diet-fed, lean *Adcy3-AdcKO* and *LoxP* control mice. We also performed indirect calorimetry assessment of energy homeostasis at 22°C and 5°C but found no differences in respiratory exchange ratios (RER, Fig.2g).

**Figure 2:**
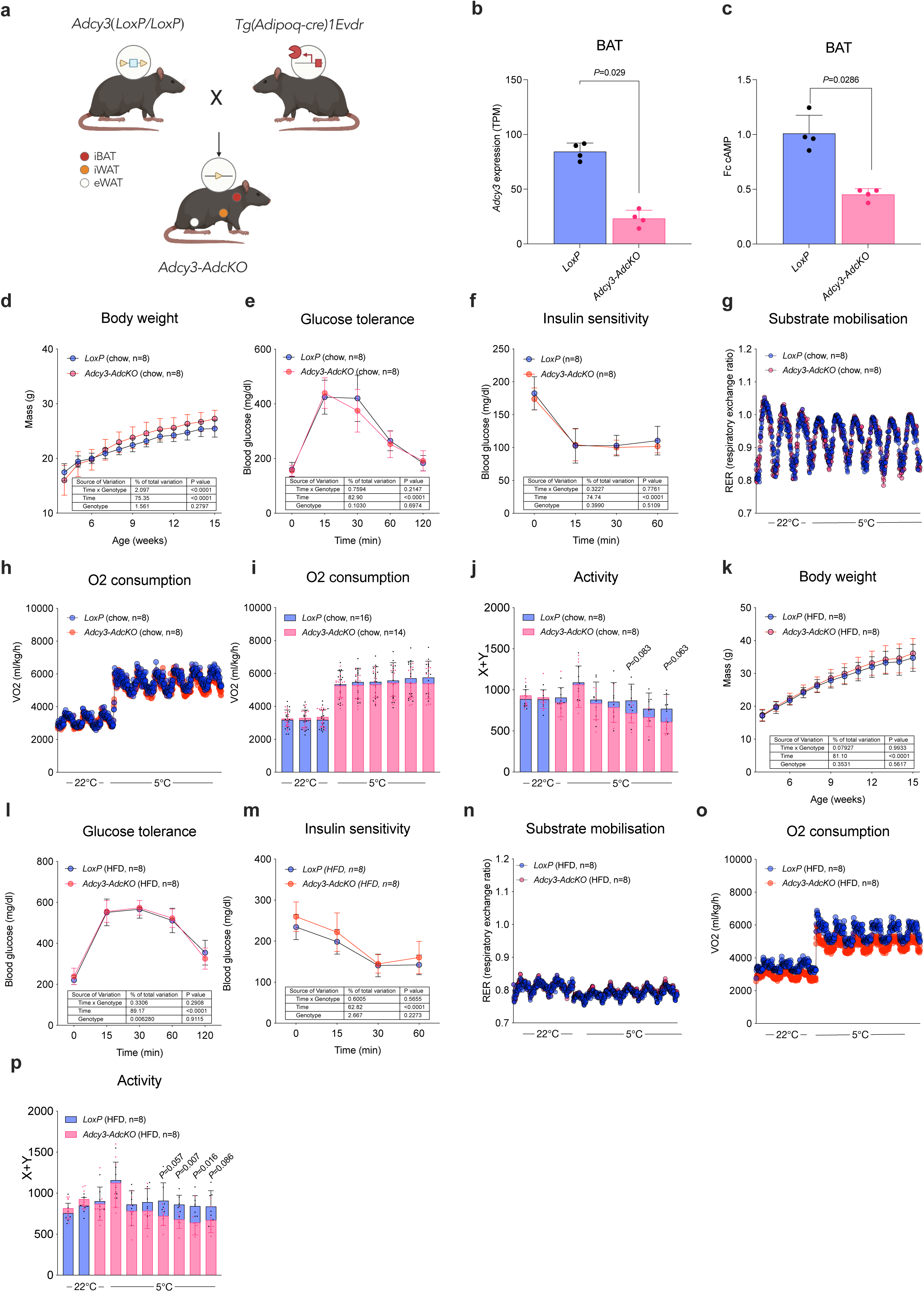
ADCY3 is required for cAMP biosynthesis in BAT and required for cold adaptation in obesity. (**a**) Illustration of breeding strategy to obtain adipocyte-specific *Adcy3* knockout mice (*Adcy3-AdcKO*) and *LoxP* controls. (**b**) Quantification of *Adcy3* expression in BAT of chow-diet fed, male *LoxP* (n=4) and *Adcy3-AdcKO* (n=4) by RNA-seq. (**c**) Determination of total cAMP levels by ELISA in BAT obtained from chow-diet fed, male *LoxP* (n=4) and *Adcy3-AdcKO* (n=4) housed at 22°C and plotted as fold-change (fc). (**b-c**) Bar graphs represent mean ± SD with all data points plotted. Unpaired, two-tailed, and non-parametric Mann-Whitney tests were performed to assess statistical significance. *P*-values are indicated within the panel. (**d**) Body weight trajectories in chow diet-fed, male *LoxP* (n=8) and *Adcy3-AdcKO* (n=8) mice. (**e-f**) Blood glucose levels during (**e**) glucose tolerance tests and (**f**) insulin tolerance tests in chow diet-fed, male *LoxP* (n=8) and *Adcy3-AdcKO* (n=8) mice. (**d-f**) Graphs represent mean ± SD with all data points plotted. Statistical significance was determined by performing Two-Way ANOVA with repeated measurements for x values (mixed models). *Post-hoc P*-value correction to account for multiple testing was performed using Bonferroni adjustment. The source of variation, % of the variation and exact *P* values are given in table insets. (**g**) Indirect calorimetric quantification of respiratory Exchange Ratios (RER) in chow-diet fed, male *LoxP* (n=8) and *Adcy3-AdcKO* (n=8) mice. (**h**) Indirect calorimetric measurement of oxygen consumption in chow-diet fed, male *LoxP* (n=8) and *Adcy3-AdcKO* (n=8) mice. (**i**) Quantification of mean daily oxygen consumption in chow-diet fed, male *LoxP* (n=16) and *Adcy3-AdcKO* (n=14) mice. (**j**) Indirect calorimetric determination of mean daily locomotor activity in chow-diet fed, male *LoxP* (n=8) and *Adcy3-AdcKO* (n=8) mice. (**i-j**) Bar graphs represent mean ± SD with all data points plotted and parametric, unpaired, two-tailed Student’s t-tests were performed to assess statistical significance. *P*-values are indicated within the panel. (**k**) BW trajectories in HFD-fed, male *LoxP* (n=8) and *Adcy3-AdcKO* (n=8) mice. (**l-m**) Blood glucose levels during (**l**) glucose tolerance tests and (**m**) insulin tolerance tests in HFD-fed, male *LoxP* (n=8) and *Adcy3-AdcKO* (n=8) mice. (**k-m**) Graphs represent mean ± SD with all data points plotted. Statistical significance was determined by performing Two-Way ANOVA with repeated measurements for x values (mixed models). *Post-hoc P*-value correction to account for multiple testing was performed using Bonferroni adjustment. The source of variation, % of the variation and exact *P* values are given in table insets. (**n**) Indirect calorimetric quantification of respiratory Exchange Ratios (RER) in HFD-fed, male *LoxP* (n=8) and *Adcy3-AdcKO* (n=8) mice. (**o**) Indirect calorimetric measurement of oxygen consumption in HFD-fed, male *LoxP* (n=8) and *Adcy3-AdcKO* (n=8) mice. (**p**) Indirect calorimetric determination of mean daily locomotor activity in HFD-fed, male *LoxP* (n=8) and *Adcy3-AdcKO* (n=8) mice. Bar graphs represent mean ± SD with all data points plotted and parametric, unpaired, two-tailed Student’s t-tests were performed to assess statistical significance. *P*-values are indicated within the panel.

In line with little overt metabolic effects following *Adcy3* loss *in vivo*, we detected no differences in regulation of genes involved in thermogenic activation in *Adcy3-AdcKO* BAT at 22°C and 5°C (Fig.S3a,b). We also investigated thermogenic activation in mature *Adcy3*-deficient BAs *in vitro* and committed *(i) Adcy3-AdcKO* stromal-vascular fraction-derived progenitor cells (SVF) or *(ii) LoxP* SVFs in mature BA, transduced *LoxP*-derived BAs with adeno-associated virus 8 (AAV8) encoding Cre (AAV8-cre), followed by CL316,243 stimulation of both models, genetic and viral. *Adcy3*-deficient and control cells showed neither changes in adipocyte identity markers like *Pparg2* and *Fabp4* nor in activation upon beta-adrenergic treatment, despite *Adcy3* silencing in both (Fig.S2c-d), suggesting unaltered BA formation and activation. Of note, we saw progressive trends towards impaired oxygen consumption (Fig.2h-i) and locomotor activity (Fig.2j), suggesting mildly impaired cold tolerance during prolonged cold exposure in *Adcy3-AdcKO* mice. Thus, adipocyte-specific loss of *Adcy3* only marginally affects thermogenesis and metabolic function in lean mice, but despite controlling cAMP levels in BAT, does not represent a major regulator for cAMP-dependent thermogenesis.

We and others reported that impediment of BAT function predisposes mice to increased body weight gain and metabolic deterioration when feeding obesogenic high-fat diets (HFD) ^37–41^. Given the modest effects of ADCY3 deficiency on energy expenditure in lean mice, we next asked if HFD renders *Adcy3-AdcKO* mice more susceptible to metabolic complications. We again observed no genotype-specific differences in BW (Fig.2k), adipose tissue weights (Fig.S3a-b), glucose (Fig.2l) or insulin tolerance (Fig.2m). When performing indirect calorimetry, obese *Adcy3-AdcKO* mice displayed a reduction in RER (Fig.2n) compared to lean mice (Fig.2g), reflecting a transition in substrate mobilization from carbohydrate to lipid oxidation in diet-induced obesity (DIO), yet no Adcy3-driven differences were seen. In contrast, oxygen consumption (Fig.2o) and activity (Fig.2p) were reduced in obese *Adcy3-AdcKO* mice compared to HFD-fed *LoxP* controls. Thus, fat-specific loss of *Adcy3* in mice reduces energy expenditure in obese mice, also during cold stress.

### ADCY3 loss causes precocious activation of PKA-independent signaling pathways

Intrigued by the lack of thermogenic and metabolic dysfunction in lean *Adcy3-AdcKO* mice, we tested whether *Adcy3* loss in adipocytes rewired cAMP signaling. To test whether kinase activities are altered in *Adcy3-AdcKO* BAT, we first performed Western Blot activity profiling using PKA substrate-specific phospho-antibodies and analyzed UCP1 protein expression. In line with lack of thermogenic dysfunction in lean mice, PKA substrate activation and UCP1 protein levels in BAT after chronic cold exposure were upregulated to same degrees in *LoxP* and *Adcy3-AdcKO* mice (Fig.S3c). To test more broadly the effects of *Adcy3* deficiency on signaling responses, we carried out serine/threonine kinase (STK) profiling using PamGene peptide arrays to globally infer differential STK kinase activities in *Adcy3-AdcKO* BAT. In *LoxP* mice, acute cold exposure (24 h) elicited only mild overall STK signaling responses (Fig.S3d), whereas chronic cold stimulation for 6 days decreased the activity of members of the CMGC kinase family (including cyclin-dependent kinases (CDK), glycogen synthase kinases (GSK) and MAP kinases (MAPK)), and activity of AGC family members (including protein kinase A (PKA), C (PKC), and G (PKG) were elevated (Fig.S3ei,ii). The CMGC family includes important hubs of cAMP-dependent signaling like the MAPKs p38 ^42^, ERK ^43^, JNK ^44^, which are typically engaged during acute thermogenic activation^45^. However, STK signatures in BAT of *Adcy3-AdcKO* at 22°C resembled chronic cold exposure in *LoxP* mice with exception that PKA activity remained unchanged (Fig.S3f, i-iii). Thus, kinase signaling pathways regulating BAT activation are indeed rewired in *Adcy3-AdcKO* mice but independent of PKA signaling. This is reflected on the functional level as minor metabolic changes were observed in lean *Adcy3-AdcKO* mice at room temperature. Only when coupled to an additional obesity burden, *Adcy3-AdcKO* are sensitive to metabolic deterioration.

### Active thermogenic adipocytes express a truncated ADCY3 (ADCY3-AT) transcript and protein isoform

Chromatin profiling by chromatin-immunoprecipitation-sequencing (ChIP-seq) allows genome-wide mapping of DNA-regulatory elements and changes in cellular energetic activation and/or differentiation are paralleled by profound remodeling of promoter (e.g., H3K4me3) and enhancer (e.g., H3K27Ac)-associated histone post-translational modifications in BA^46–48^. When interrogating previously published H3K4me3 data from BAT of cold-exposed mice^49^, we observed a H3K4me3-marked promoter in intron 2 of the *Adcy3* gene that arose during cold (Fig.S4a), which we verified by ChIP-qPCR (Fig.S4b). This promoter marked a cold-dependent novel transcriptional start site giving rise to a 5’-truncated *Adcy3* mRNA isoform (Fig.S4a), which we termed *Adcy3-at* to discern it from GENCODE annotated full-length *Adcy3* (*Adcy3-fl*). When combining Illumina short-read with Oxford Nanopore Technology full-length RNA-seq from cold-activated BAT, we confirmed that *Adcy3-at* represents a contiguous transcript ranging from *Adcy3-at* TSS in exon 2b to exon 22 (Fig.3a). *Adcy3-at* encoded ADCY3-AT proteins are predicted as N-terminally truncated compared to full-length ADCY-FL proteins (Fig.3b) and distinct from ADCY3 loss-of-function protein isoforms previously reported in human populations ^17–19^. Thus, cold exposure activates an intronic *Adcy3* promoter that produces N-terminally truncated, unannotated *Adcy3-at* mRNA isoform in thermogenic fat depots.

**Figure 3:**
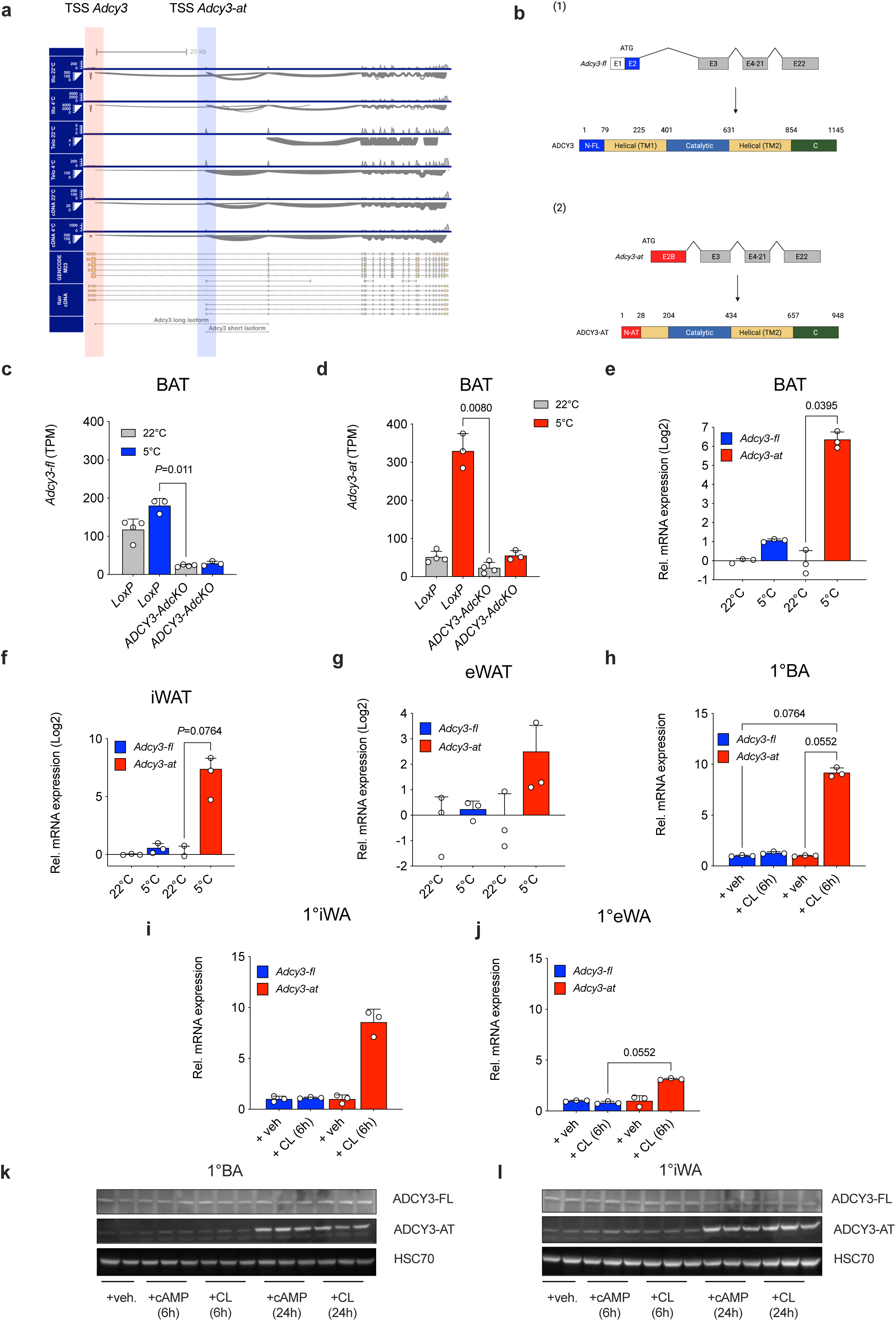
Activated brown adipocytes express a truncated ADCY3 (ADCY3-AT) transcript and protein isoform. (**a**) Sashimi plots visualizing splicing junctions from aligned RNA-seq data in BAT in the murine *Adcy3* gene from 22°C-housed 20 weeks old male wildtype C57BL/6N mice or from animals singly housed at 4 °C for a period of 24 h prior to harvesting adipose tissues. Data represent n=3-5 individual mice per condition. *Illu*, Illumina short-read RNA-seq; *Telo*, TeloPrime Full-Length cDNA-seq; *cDNA*, Direct cDNA-seq. Reads were aligned against GENCODE M29 annotation and transcript re-assembly using Illumina short-read and full-length RNA-seq using FLAIR^75^. (**b**) Schematic illustration of (1) full-length *Adcy3-fl* transcript and ADCY3-FL protein structure, and (2) *Adcy3-at* transcript and ADCY3-AT protein structure (2). (**c-d**) Expression of *Adcy3-fl* (**c**) and *Adcy3-at* (**d**) in BAT of chow diet-fed animals housed at 22°C (n=4 *LoxP*, n=4 *Adcy3-AdcKO*) and 5°C (n=3 for *LoxP*, n=3 for *Adcy3-AdcKO*) determined by RNA-seq. (**e-g**) Relative expression of *Adcy3-fl* and *Adcy3-at* as determined by quantitative PCR analysis of BAT (**e**), iWAT (**f**) and eWAT (**g**) of male C57BL/6N mice housed at room temperature (n=3) or exposed to 5°C for 24 h (n=3). (**h-j**) Relative expression of *Adcy3-fl* and *Adcy3-at* as determined by quantitative PCR analysis of primary adipocytes derived from stromal-vascular fraction (SVF) cells isolated from BAT (**h**), iWAT (**i**), and eWAT (**j**) depots and stimulated with 6 h of 10 µM CL316,243. Replicates represent primary adipocytes isolated from individual mice (n=3 mice per depot and stimulation regimen). (**c-j**) Bar graphs represent mean ± SD with all data points plotted. To test for statistical significance, non-parametric (ranked) Kruskal–Wallis one-way analysis of variance (ANOVA) tests with Dunn’s correction for multiple testing were performed. *P*-values are indicated in the panel. (**k-l**) Immunoblot analysis of primary adipocytes differentiated from SVF cells isolated from BAT (1°BA) and iWAT (1°iWA), stimulated for 6 h and 24 h with 1 mM db-cAMP (cAMP) and 10 µM CL316,243 (CL) and blotted using pan anti-ADCY3 antibody. Bands corresponding to ADCY3-FL and ADCY3-AT are from the same membrane but represent diffent exposure times. Calnexin (CLNX) was used as loading control.

When analyzing our RNA-seq data separately for *Adcy3-at* and *Adcy3-fl* isoform expression, we found that predominantly *Adcy3-at* was induced during cold exposure and constituted the dominant *Adcy3* isoform after cold as shown by RNA-seq (Fig.3c-d) and qPCR (Fig.3e) quantification. Other thermogenic depots like iWAT also induced *Adcy3-at* in cold (Fig.3f), whilst eWAT *Adcy3-at* induction was not significant (Fig.3g), pointing to roles of ADCY3-AT in thermogenic processes in mature BA, yet effects of ADCY3-AT on brown adipogenesis were not addressed at point. To address this important point specifically, we analyzed *Adcy3* isoform expression during adipogenic differentiation and observed that *Adcy3-at* expression was low throughout adipogenesis of BAT SVFs into BA and was only upregulated upon dibutyryl (db)-cAMP and CL316,243 stimulation (Fig.S4e), whereas *Ucp1* and *Adcy3-fl* expression was more highly expressed during brown adipogenesis (Fig.S4f-g). *Adcy3-at* was upregulated in a cell-autonomous manner in primary brown adipocytes (1°BA) and primary inguinal white adipocytes (1°iWA), yet only trended towards induction in primary epididymal white adipocytes (1°eWA) after stimulation with beta3-adrenergic agonist CL316,243 in all depots (Fig.3h-j). *Adcy3-fl*, but especially *Adcy3-at*, were expressed at low abundance in non-adipose metabolic tissues like liver, skeletal muscle, kidney, and pancreas (Fig.S4c-d). In obesity- and aging-associated adipose dysfunction, *Adcy3-at* expression was unchanged in BAT and downregulated in iWAT of DIO mice, a finding seen for canonical BAT markers like *Ucp1*, Iodothyronine Deiodinase 2 (*Dio2*) and ELOVL fatty acid elongase 3 (*Elovl3*, Fig.S4h,i). *Adcy3-at* expression was also unaffected in BAT of aged (25 months-old) compared to young (2 months-old) C57BL/6N male mice (Fig.S4j). Thus, two mouse models of functional thermogenic decline ^50^ showed no differences in BAT *Adcy3-at* expression.

The ADCY3-AT and ADCY3-FL amino acid (aa) sequences are for large parts identical but differ at the N-terminus: ADCY3-AT lacks 146aa and instead contains a novel 28aa N-terminus, entailing that ADCY3-AT proteins specifically lack the 1^st^ block of transmembrane (TM) domains, which constitutes an integral structural hallmark and functional feature of most adenylyl cyclases (Fig.3b). When performing Western blot analysis using a C-terminal pan-ADCY3 antibody, we detected an additional ADCY3-specific signal at ca. 70 kD after 24 h of db-cAMP and CL316,243 stimulation in 1°BA and 1°WA, which was absent in unstimulated cells (Fig.3k-l), likely representing ADCY3-AT proteins. Thus, pharmacological and cold temperature-mediated activation of thermogenic adipocytes leads to presence of a novel intronic *Adcy3-at* promoter that encodes truncated ADCY3-AT proteins.

### ADCY3-AT inhibits oxidative metabolism in vitro and in vivo

As *Adcy3-AdcKO* mutants were deficient for both *Adcy3* isoforms (Fig.3c-d), we tested the specific role of ADCY3-AT in cellular and metabolic homeostasis of BA *in vivo* and *in vitro.* We performed CRISPR-Cas9-mediated gene deletion using single-guide RNAs (sgRNAs) flanking the H3K4me3-marked *Adcy3-at* promoter and excised a 978 bp-long DNA fragment including the *Adcy3-at* TSS in embryonic stem cells (Fig.S5a). Excision of *Adcy3-at* TSS and promoter regions was confirmed by Southern blot (Fig.S5b) and genomic PCR analysis (Fig.S5c). Homozygous ADCY3-AT null animals, henceforth termed *Adcy3ΔAT*, exhibited no overt phenotypes, were fertile in both sexes, and *Adcy3ΔAT* alleles were inherited at Mendelian ratios, arguing against confounding developmental defects after *Adcy3-at* deletion. Importantly, *Adcy3ΔAT* mice were devoid of *Adcy3-at* (Fig.4b), but not *Adcy3-fl* (Fig.4c).

**Figure 4:**
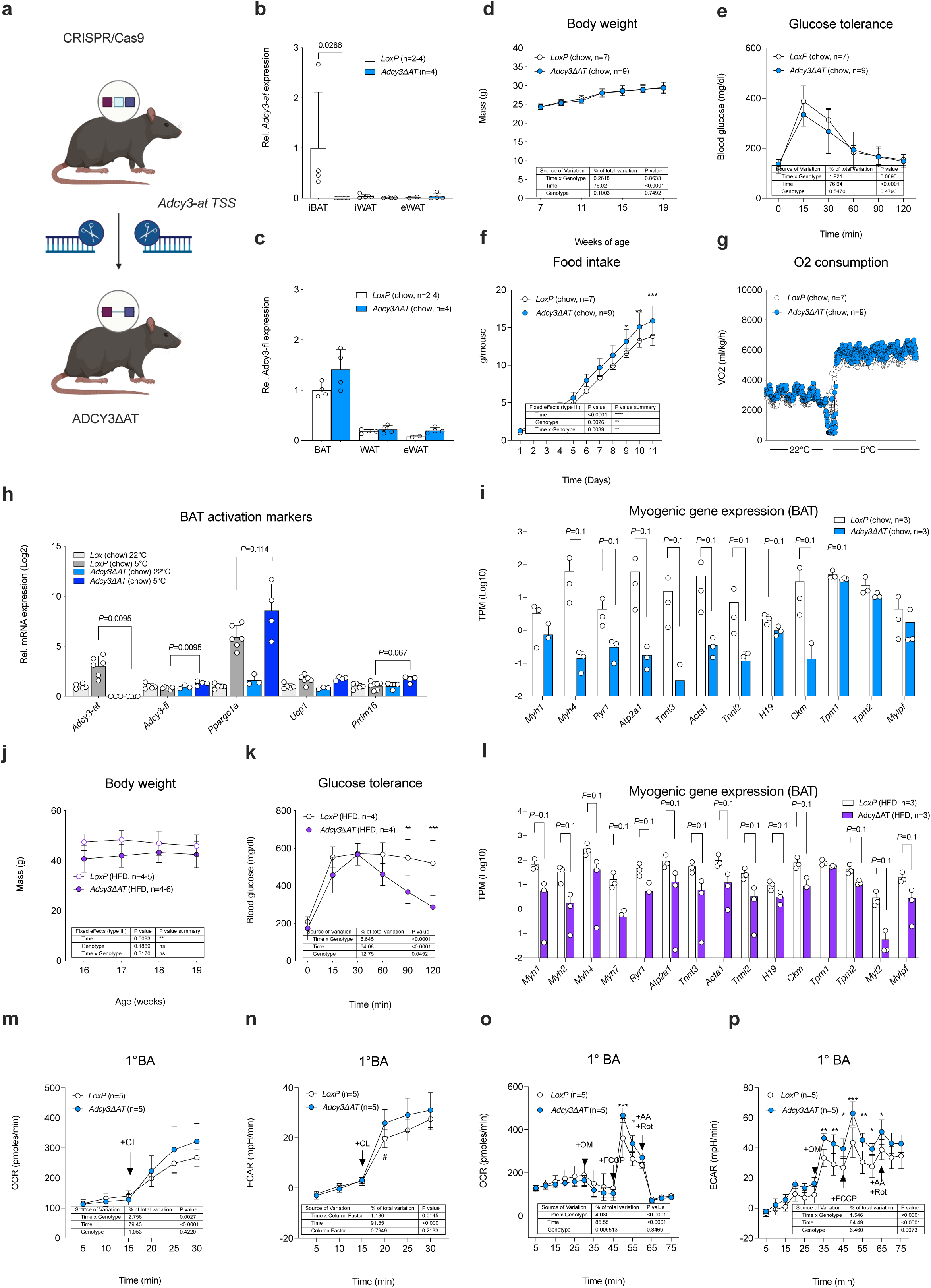
ADCY3-AT inhibits oxidative metabolism in vitro and in vivo. (**a**) Illustration of CRISPR-Cas9 mediated excision of *Adcy3-at* specific exon 2b including *Adcy3-at* TSS (light blue) yielding ADCY3-AT specific knockout mice. (**b-c**) Relative expression of (**b**) *Adcy3-at* and (**c**) *Adcy3-fl* determined by qPCR analysis of BAT, iWAT, and eWAT of chow diet fed, male *LoxP* and *Adcy3ΔAT* mice housed at 22°C (n=3-4 per tissue and genotype). Bar graphs represent mean ± SD with all data points plotted. An unpaired, two-tailed, and non-parametric Mann-Whitney tests were performed to assess statistical significance. *P*-values are indicated within the panel. (**d**) Body weights in chow diet-fed, male *LoxP* (n=7) and *Adcy3ΔAT* (n=9) mice. (**e**) Blood glucose excursion during glucose tolerance test in chow diet-fed, male *LoxP* (n=7) and *Adcy3ΔAT* (n=9) mice. (**f**) Cumulative food intake as measured by indirect calorimetry in chow diet-fed, male *LoxP* (n=7) and *Adcy3ΔAT* (n=9) mice. (**g**) Indirect calorimetric measurement of oxygen consumption in chow-diet fed, male *LoxP* (n=7) and *Adcy3ΔAT* (n=9) mice. (**d-g**) Graphs represent mean ± SD with all data points plotted. Statistical significance was determined by performing Two-Way ANOVA with repeated measurements for x values (mixed models). *Post-hoc P*-value correction to account for multiple testing was performed using Bonferroni adjustment. The source of variation, % of the variation and exact *P* values are given in table insets. (**h-i,l**) Relative expression of indicated mRNAs in (**h**) chow diet-fed, male *LoxP* (n=5 for 22°C, n=6 for 5°C) and *Adcy3ΔAT* (n=3 for 22°C, n=4 for 5°C), (**i**) chow diet fed, male *LoxP* (n=3) and *Adcy3ΔAT* (n=3) and (**l**) HFD-fed, male *LoxP* (n=3) and *Adcy3ΔAT* (n=3) mice. Bar graphs represent mean ± SD with all data points plotted and unpaired, two-tailed, non-parametric Mann-Whitney tests were performed for indicated comparisons to assess statistical significance. *P*-values are given within the panel. (**j**) BW in 16 to 19 weeks old, HFD-fed male *LoxP* (n=4-5) and *Adcy3ΔAT* (n=4-6) mice. (**k**) Blood glucose during glucose tolerance tests in HFD-fed, male *LoxP* (n=4) and *Adcy3ΔAT* (n=4) mice. (**j-k**) Graphs represent mean ± SD with all data points plotted. Statistical significance was determined by performing Two-Way ANOVA with repeated measurements for x values (mixed models). *Post-hoc P*-value correction to account for multiple testing was performed using Bonferroni adjustment. The source of variation, % of the variation and exact *P* values are given in the table insets. (**m,n**) Oxygen consumption rates (**m**) and extracellular acidification rates (**n**) in 1°BA derived from SVF of *LoxP* or *Adcy3ΔAT* mice after stimulation with 10 µM CL316,243. The numbers of measured wells are indicated in the panel. (**o-p**) Oxygen consumption rates (**o**) and extracellular acidification rates (**p**) in 1°BA derived from SVF of *LoxP* or *Adcy3ΔAT* mice. OM, oligomycin; FCCP, AA, Antimycin A; Rot, Rotenone. The numbers of measured wells are indicated in the panel. (**m-o**) Graphs represent mean ± SD with all data points plotted. Statistical significance was determined by performing Two-Way ANOVA with repeated measurements for x values (mixed models). *Post-hoc P*-value correction to account for multiple testing was performed using Bonferroni adjustment. The source of variation, % of the variation and exact *P* values are given in the table insets.

First, we tested the role of ADCY3-AT in regulating energy homeostasis in lean mice: Although BW remained unchanged (Fig.4d), *Adcy3ΔAT* animals exhibited improved glucose tolerance (Fig.4e) despite even mildly increased food intake (Fig.4f), normal insulin sensitivity (Fig.S5d) and unaltered locomotor activity (Fig.S5e), suggesting negative energy balance in *ADCY3ΔAT* mice. RER quotients unveiled that *Adcy3ΔAT* animals trended towards increased carbohydrate usage at 22°C (Fig.S5f), likely reflecting an increased capacity of *ADCY3ΔAT* mice for enhanced glucose disposal, which consequently exhibited trends towards increased oxygen consumption during 22°C and 5°C ambient temperatures (Fig.4g). To test for BAT-specific involvement in the improved metabolism seen in *Adcy3ΔAT* mice, we performed qPCR analysis and observed trends towards increased *Ppargc1a* and *Prdm16* expression, suggesting mildly elevated activation of cold-acclimatized *Adcy3ΔAT* BAT (Fig.4h). Notably, expression of gene markers reflecting myogenic-to-BA remodeling, such as myosin heavy chain 1 *(Myh1)* and 4 (*Myh4*), ryanodine receptor 1 *(Ryr1)*, ATPase sarcoplasmic/endoplasmic reticulum Ca2+ transporting 1 *(Atp2a1)*, troponin T3 fast skeletal type *(Tnnt3)*, actin alpha 1, skeletal muscle *(Acta1)*, troponin I2, fast skeletal type *(Tnni2)*, H19 imprinted maternally expressed transcript *(H19)*, creatine kinase, M-type *(Ckm)*, tropomyosin 1 and 2 (*Tpm1,2*) were repressed (Fig.4i), reflecting transcriptional signatures reminiscent of activated BA at the expense of myogenic gene expression in *Adcy3ΔAT* mice, a change in transcriptional signatures reported for other models of BAT gain-of-function ^38, 40^.

Albeit these mild increases in energy dissipation, we hypothesized that *Adcy3ΔAT* mice are protected against DIO-induced metabolic alterations. To test this, we exposed *LoxP* and *Adcy3ΔAT* to HFD feeding and observed that *Adcy3ΔAT* were indeed protected against DIO-associated BW gains and glucose intolerance (Fig.4j-k) and *Adcy3ΔAT* mice exhibited lower glucose levels during an insulin challenge (Fig.S5h). As seen in chow diet-fed animals, obese *Adcy3ΔAT* exhibited decreased myogenic gene expression reflecting increased BA activation (Fig.4l). To test for BA-intrinsic roles of ADCY3-AT, we performed Seahorse bioenergetic analyses in 1°BA and 1°iWA and observed elevated maximal oxygen consumption rates after CL316,243 stimulation (Fig.4m) and during FCCP treatment in mitochondrial stress tests in *Adcy3-at* deficient adipocytes (Fig.4o). Together with higher extracellular acidification rates in *Adcy3ΔAT* cells (Fig.4n,p), the results indicated a combination of increased oxidative phosphorylation and glucose oxidation in *Adcy3ΔAT* 1°BA (Fig.4m-p) and 1°WA (Fig.S5i-l). Thus, ADCY3-AT deficiency improves metabolism *in vivo*, especially upon diet-induced obesity, by increasing energy dissipation in brown adipocytes and myogenic-to-BA transcriptional remodeling.

### ADCY3-AT alters ADCY3-FL subcellular localization and thereby limits cAMP biosynthesis

Our *in vitro* and *in vivo* data thus far indicated that silencing of cold-inducible ADCY3-AT increases oxidative metabolism and thermogenic activation of BA. To test this molecularly, we analyzed STK activities using PamGene Array analysis in BAT of *Adcy3ΔAT*. At 22°C, STK activity in *Adcy3ΔAT* BAT remained largely unchanged compared to BAT from *LoxP* control mice (Fig.S6a), demonstrating that ADCY3-AT does not profoundly affect STK kinase activity under basal conditions. Upon chronic cold stimulation, directional changes in kinase activities in *Adcy3ΔAT* were comparable also to cold-exposed *LoxP* mice (Fig.S6b). Importantly, when comparing kinase activity in BAT from *Adcy3ΔAT* with *Adcy3-AdcKO*, signaling inversely correlated, indicating that deleting both ADCY3 proteins evokes opposing effects to ablating ADCY3-AT alone (Fig.S6c-d).

The presence of ADCY3-AT coincides with ADCY3-FL in activated BAT (Fig.3b,c), yet the functional implication of this co-expression remained elusive. Adenylyl cyclases are active as protein dimers, and dimerization affects protein transport to the plasma membrane (PM). Besides homodimerization, ADCYs can also assemble in heterodimers between different adenylyl cyclase isoforms^51^. We thus hypothesized that ADCY3-AT controls the activity of ADCY3-FL by protein interaction. To test this notion *in vitro*, we co-expressed FLAG-tagged ADCY3-AT and HA-tagged ADCY3-FL in Chinese Hamster Ovary (CHO) cells and demonstrated by co-immunoprecipitation that ADCY3-AT and ADCY3-FL indeed interact (Fig.5a-b). The N terminus of adenylyl cyclases control transport to PM, thereby regulating its localization and, concomitantly, enzymatic function ^52^. As ADCY3-AT lacks the N-terminal domain and interacts with ADCY3-FL, we next tested if ADCY3-AT might not be able to localize to the PM, and in turn, also impairs membrane transport of ADCY3-FL. To this end, we performed biotinylation assays to selectively labeled TM proteins that face the extracellular *environment*. When individually overexpressing the two different isoforms, only ADCY3-FL, but not ADCY3-AT, was biotinylated (Fig.5c-d), demonstrating that ADCY3-AT does not localize to the PM. To complement this result with image-based approaches, we subjected CHO cells grown on glass coverslips to short ultrasound pulses, whereby the upper part of cells was removed and the PM and its associated complexes remained intact as membrane sheets (‘unroofing’)^53^. As positive control for visualizing the PM, we expressed membrane-anchored mCherry. ADCY3-FL colocalized with mCherry in the PM, but showed little overlap with calnexin, a protein residing in the ER membrane and here specifically marking ER-PM contact sites. In contrast, ADCY3-AT showed no colocalization with mCherry, but colocalized with calnexin, demonstrating that ADCY3-AT is retained in the ER (Fig.5e-f, Fig.S6e-f). Upon co-expression of ADCY3-AT and ADCY3-FL, ADCY3-FL was retained in the ER and no longer localized to the PM, indicating that interaction of ADCY3-AT with ADCY3-FL prevented PM trafficking of ADCY3-FL proteins (Fig.5g, Fig.S6g). To probe if ADCY3-AT intrinsically possesses cAMP catalytic activity, we determined cAMP levels by ELISA in CHO cells overexpressing the individual ADCY3 isoforms. As expected, ADCY3-FL, but not ADCY3-AT, activity was stimulated with forskolin, a direct ADCY activator (Fig.S6h), and isoproterenol, a beta-adrenergic receptor agonist, which stimulates adenylyl cyclase activity via G-proteins (Fig.S6i), indicating that ADCY3-AT does not possess enzymatic activity or lost its interaction with G-proteins due to altered subcellular localization. Our hypothesis that ADCY3-AT/ADCY3-FL heterodimer formation is favored in cold-activated BAT was supported by *in silico* modeling of ADCY3-AT and ADCY3-FL heteromeric structures using AlphaFold ^54^. Our structural modelling suggested that ADCY3 heterodimer are energetically plausible (Fig.5h), underlining that ADCY3-AT-mediated retention of ADCY3-FL represents a credible scenario for ADCY3-AT-mediated repression of adenylyl cyclase activity. Thus, in the presence of both isoforms, ADCY3-AT and ADCY3-FL, ADCY3-FL transport to PM in ADCY3-AT/FL heterodimrs is impaired, suggesting that interaction of ADCY3-AT with ADCY3-FL curbs cytoplasmic cAMP production by altered localization and/or cAMP compartmentalization.

**Figure 5:**
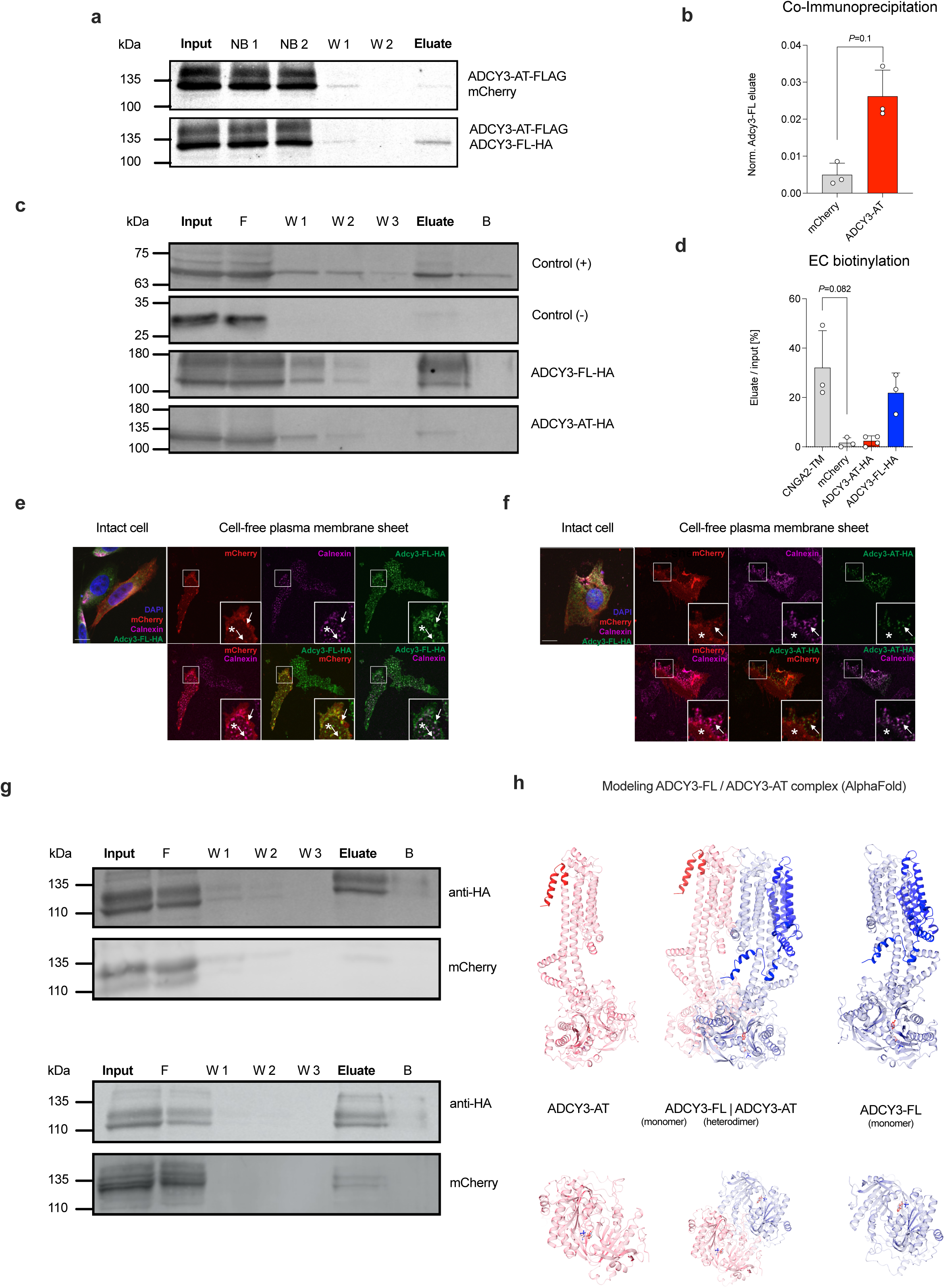
ADCY3-AT alters ADCY3 subcellular localization and thereby limits cAMP biosynthesis. (**a**) Co-immunoprecipitation of ADCY3-AT-FLAG and ADCY3-FL-HA in CHO cells. CHO K1 cells stably expressing ADCY3-FL-HA were transiently transfected with mCherry (upper row) or ADCY3-AT-Flag (bottom row). Total protein lysates were incubated with anti-FLAG magnetic beads and the different fractions were analyzed by Western blot (Input, NB: non-bound, W: wash, Eluate). (**b**) Quantification of Western blot analyses. The ratio of ADCY3-FL-HA protein density in the eluate of total protein lysates described in (**a**). Bar graphs represent mean ± SD with all data points plotted. An unpaired, two-tailed, and non-parametric Mann-Whitney tests were performed to assess statistical significance. If relevant, *P*-values are indicated within the panel. (**c**) Biotinylation assay. Representative Western blot analysis of total protein lysates isolated from CHO cells stably expressing CNGA2-TM (+, CNB-3B10), transiently transfected with mCherry (-, anti-RFP antibody), or stably expressing ADCY3-FL-HA or ADCY3-AT-HA (anti-HA antibody). Cells were treated with the biotinylation reagent Sulfo-NHS-SS-Biotin, lysed (Input), and purified using a NeutrAvidin-agarose-resin (F: flow through, W: wash, E: eluate, B: beads). (**d**) Quantification biotinylation assay. Densitometric analysis of n=3 (CNGA2-TM, mCherry, ADCY3-FL-HA) and n=4 (ADCY3-AT-HA) Western blots shown in (**e**). Values were normalized to the respective input sample. The ratio of eluate to input in % is shown as mean ± SD with all data points plotted. To test for statistical significance, non-parametric (ranked) Kruskal– Wallis one-way analysis of variance (ANOVA) tests with Dunn’s correction for multiple testing were performed. *P*-values are indicated in the panel. (**e**) CHO cells stably expressing ADCY3-FL-HA and transiently transfected with mCherry-CAAX were sonicated and labeled with an anti-Calnexin antibody (ER-marker, magenta) and an anti-HA antibody (green). mCherryCAAX fluorescence is indicated in red. Single channels are shown at the top, merged images at the bottom row. White boxes are shown as magnified view at the bottom-right. Arrows point to endoplasmic reticulum-plasma membrane (ER-PM) contact sites, asterixis indicate contact-free parts within the membrane sheet. Scale bar: 10 μm. (**f**) See (**e**) for CHO cells stably expressing ADCY3-AT-HA. (**g**) Representative Western blot analysis of total protein lysates isolated from CHO K1 cells stably expressing ADCY3-FL-HA or, transiently expressing (*top*) mCherry or (*bottom*) ADCY3-AT-FLAG-2A-mCherry. Cells were treated with the biotinylation reagent Sulfo-NHS-SS-Biotin and the protein lysate was purified using a NeutrAvidin-agarose-resin. The following samples were loaded: Input, Flow-through (F), Wash ( W1-3), Eluate, Beads (B). The presence of Adcy3-FL-HA in the plasma membrane was verified using an anti-HA antibody, an anti-mCherry antibody was used as a control. (**h**) AlphaFold modeling of ADCY3-FL/ADCY3-AT heterodimer complex. Full-colored and transparent domains depict non-conserved and conserved residues between ADCY3-FL and ADCY3-AT, respectively.

### Conserved and cold-inducible PPARGC1A -AT drives Adcy3-at expression in brown adipocytes

PPARGC1A represents a central regulator of mitochondrial biogenesis ^29, 55^ and is imperative for brown adipocyte differentiation and function ^56^. Intriguingly, alternative splicing-mediated removal of exons 6-7 of *Ppargc1a* has been reported to results in C-terminally truncated NT-PGC1alpha ^57–59^. Loss of NT-PGC1alpha protects against obesity ^60^, suggesting negative roles for truncated PPARGC1A protein isoforms in BA differentiation and/or function. Beyond NT-PGC1alpha, alternative promoter usage and alternative splicing events give rise to more than ten PPARGC1A proteoforms, each coordinating tissue- and state-dependent adaptive transcriptional processes ^29, 61^. When interrogating our Illumina and Nanopore RNA-seq data, we observed the cold-induced alternative promoter accrual and generation of a novel transcript start site 5’ of the canonical, full-length *Ppargc1a* (*Ppargc1a-fl*, i.e. the *Pgc1alpha1* isoforms reported previously^61^) TSS, giving rise to a novel *Ppargc1a* mRNA isoform. In analogy to *Adcy3-at* we termed this 5’-terminally truncated mRNA isoform *Ppargc1a-at* (Fig.6a). *Ppargc1a-at* encoded a C-terminally truncated PPARGC1A-AT proteoform (Fig.6b) that represents a murine homologue to the *Ppargc1alpha4* isoform reported for human skeletal muscle ^28^. As PPARGC1A, together with cognate nuclear hormone transcription factors, controls adaptive gene expression programs ^62^, we postulated that induction of PPARGC1A-AT could be implicated in cold-adaptive BA responses involving *Adcy3-at*. Akin to *Adcy3-at*, *Ppargc1a-at* was only induced in thermogenic adipose depots during cold, whereas *Ppargc1a-fl* remained unchanged in all depots investigated (Fig.6c). To test the specific implication of PPARGC1A-AT in *Adcy3-at* expression, we designed two locked nucleic acid inhibitors to silence *(i) Ppargc1a-fl* and *Ppargc1a-at* in combination or *(ii) Ppargc1a-at* in 1°BA. Noteworthy, knocking down *Ppargc1a* did not affect *Adcy3-fl*, but prevented CL316,243 mediated *Adcy3-at* induction (Fig.6d-e), illustrating that PPARGC1A, likely by virtue of truncated PPARGC1A-AT is implicated in *Adcy3-at* transcriptional regulation.

**Figure 6:**
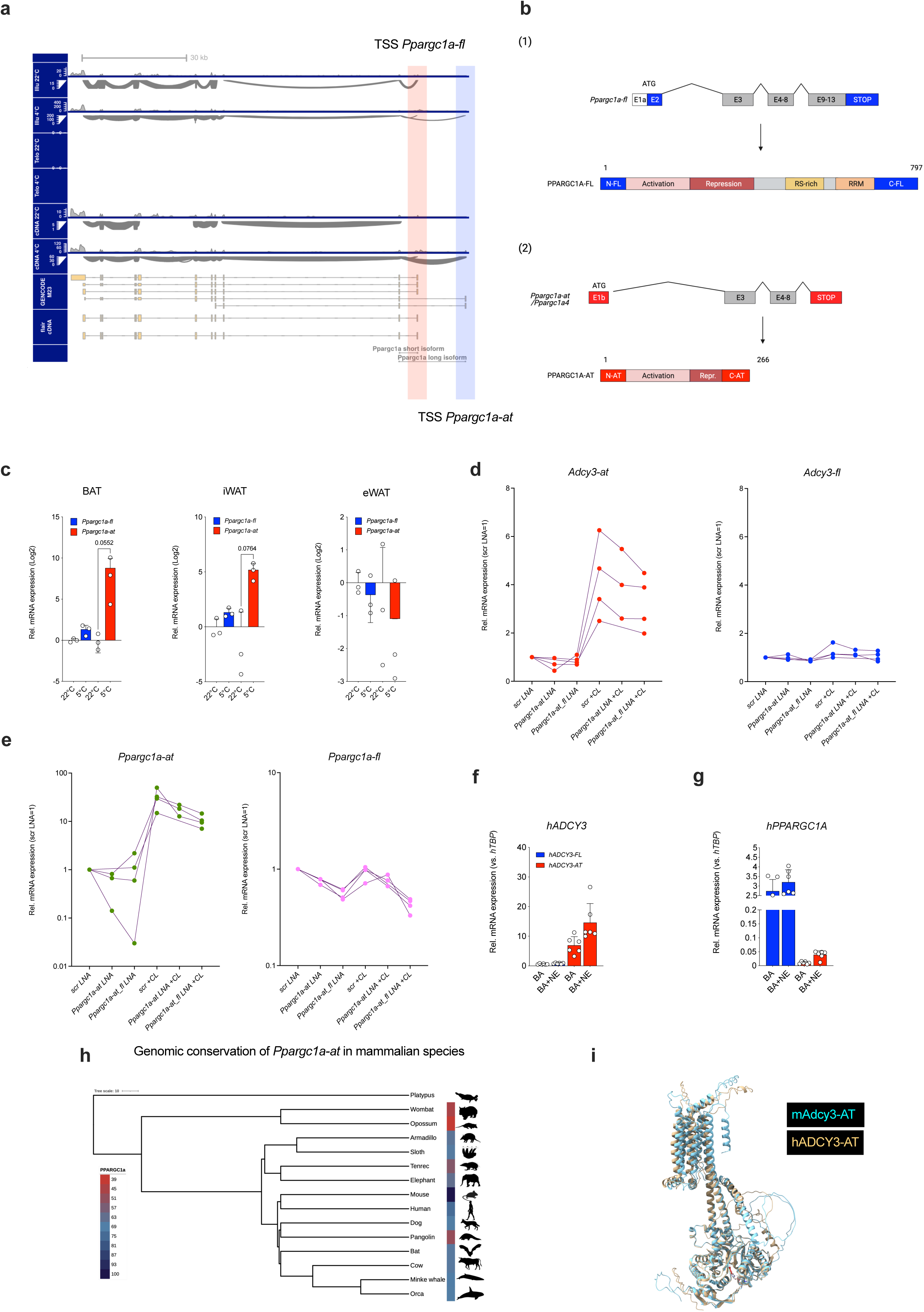
Conserved and cold-inducible PPARGC1A-AT drives *Adcy3-at* expression in brown adipocytes. **(a)** Sashimi plots visualizing splicing junctions from aligned RNA-seq data in BAT in the murine *Ppargc1a* gene locus from 22°C-housed 20 weeks old male wildtype C57BL/6N mice or animals singly housed at 4 °C for a period of 24 h prior to harvesting adipose tissue. n=3-5 animals per temperature. *Illu*, Illumina short-read RNA-seq; *Telo*, TeloPrime Full-Length cDNA-seq; *cDNA*, Direct cDNA-seq. Reads were aligned against GENCODE M29 annotation and transcript re-assembly using Illumina and ONT information using FLAIR^75^. (**b**) Illustration of full-length *Ppargc1a-fl* transcript and PPARGC1A-FL protein structure (1) as well as *Ppargc1a-at* transcript and PPARGC1A-AT protein structure (2). (**c**) Relative expression of *Ppargc1a-fl* and *Ppargc1a-at* as determined by quantitative PCR analysis of primary adipocytes derived from stromal-vascular fraction (SVF) cells isolated from BAT (*left*), iWAT (*middle*), and eWAT (*right*) depots. Replicates represent primary adipocytes isolated from individual mice (n=3 mice per adipose depot and stimulation regimen). Bar graphs represent mean ± SD with all data points plotted. To test for statistical significance, non-parametric (ranked) Kruskal–Wallis one-way analysis of variance (ANOVA) tests with Dunn’s correction for multiple testing were performed. *P*-values are indicated in the panel. (**d-e***) Adcy3* (**d**) and *Ppargc1a* (**e**) isoform expression in 1°BA after transfection with 25 nM scrambled (scr LNA) or *Ppargc1a* locked nucleic acids (LNAs) targeting both *Ppargc1a* (*Ppargc1a-at_fl* LNA) isoforms or Ppargc1a-at exclusively (*Ppargc1a-at* LNA) knockdown. Data represent three to four independent experiments, each performed in three technical replicates. Paired samples are represented by individual lines and scr LNA set to unity =1. Line graphs represent mean ± SD with all data points plotted and paired, two-tailed Student’s T-tests were performed to assess statistical significance. *P*-values are indicated within the panel. (**f-i**) Expression of human *ADCY3-AT* (**f**) and *PPARGC1A-AT* (**g**) in primary human brown adipocytes (hBA) derived from stromal-vascular precursors and stimulated with 1 µM norepinephrine (NE) for 16 h as described previously^76^. (**h**) Genomic conservation of *Ppargc1a-at* among representative mammals. Color gradient indicates percent nucleotide identity of exon 1b relative to mouse. Phylogeny based on ^77^. (**i**) Overlap of predicted structure (AlphaFold) of mouse and human ADCY3-AT proteins. Cyan and Gold depict mouse and human isoform, respectively

Finally, to test if cold-inducible ADCY3/PPARGC1A alternative promoter accrual events are conserved in humans, we isolated SVF from deep-neck biopsies of human individuals, differentiated these into human white and brown adipocytes and performed qPCR for full-length and truncated *Ppargc1a* and *Adcy3* transcripts after norepinephrine (NE) stimulation. We observed NE-induced *ADCY3-AT* and *PPARGC1A-AT* expression in brown human adipocytes (Fig.6f,g). We also investigated the broader evolutionary context of *Ppargc1a-at* and *Adcy3-at* by sequence comparison across a panel of thermogenic and non-thermogenic species (Fig.S7c). We found that *Ppargc1a-at* exon 1 was broadly conserved across eutherian species (Fig.6j, Fig.S7d,e), whereas mouse *Adcy3-at* was only conserved within selected rodent species (Fig.S7f) and likely arose independently from human *ADCY3-AT* that was only conserved in primates (Fig.S7g). Despite this convergent ADCY3-AT evolution, rodent and primate *Adcy3-at* transcripts encoded sequence- and structurally related proteins (Fig.6k). Thus, cold stress employs an evolutionarily ancient signal transduction axis that couples truncated PPARGC1A-AT proteins to cold-induced, rheostatic transcriptional responses in activated brown adipocyte that, amongst others, involves ADCY3-AT.

## Discussion

Since the re-discovery of active and recruitable BAT in humans^3–5^, enhancing brown fat activity has been increasingly recognized as innovative therapeutic regimen to mitigate obesity and obesity-related health decline. Understanding BAT activity and increasing energy expenditure by ectopic BAT activation can thus become key to control metabolic health in a positive manner. Whilst the molecular circuitry underlying thermogenic activation of brown fat is becoming increasingly well understood, we lack detailed understanding of how BAT activation is kept in balance to avoid negative consequences of exaggerated energy dissipation. Here, we have demonstrated a novel, evolutionary conserved molecular mechanism that functions as a rheostat to maintain BAT function. This is based on the expression of an N-terminally truncated ADCY3-AT isoform that regulates cAMP synthesis during chronic cold exposure and, thereby, maintains BAT energy homeostasis.

Expression of novel ADCY isoforms has also been demonstrated in other cell types, i.e., vascular smooth muscle cells (VSMCs). A crucial change in VSMC function is the trans-differentiation from a from contractile/quiescent to a secretory/proliferative phenotype, which underlies artherogenesis and vascular remodeling ^63^. Trans-differentiation in response to the pro-atherogenic cytokine interleukin-1beta (IL-1b) relies on *de novo* expression of ADCY8 in VSMCs ^64, 65^. However, an increase in intracellular cAMP levels is supposed to inhibit the secretory/proliferative VSCM phenotype ^66, 67^. Strikingly, the ADCY8 isoforms expressed in VSCMs upon IL-1b stimulation are splice variants which results in a shorter ADCY8 proteins, which are catalytically inactive. N-terminally truncated ADCY8 proteins act in a dominant-negative manner on other ADCY isoforms by forming heterodimers, retaining them in the ER and reducing intracellular cAMP levels ^68, 69^. This is in line with our results, mirroring our observation of a stimulus-dependent *de novo* expression of truncated ADCY3-AT in BAT. Thus, we here delineate that balancing cAMP levels by expressing truncated ADCY isoforms is a more broadly applicable mechanisms that occurs not only in VSCMs but also in thermogenic fat and across different ADCY isoforms. Whether the regulation of *de novo* expression is regulated by common signaling pathways in different cell types, is not known. We here identified a H3K4me3-marked promoter in intron 2 of the *Adcy3* gene, selectively induced upon cold, resulting in novel transcriptional start site for a 5’-truncated *Adcy3* mRNA isoform. The regulation of epigenetic modifications in the context of cAMP signaling is unknown, but it has been shown that the expression of a histone demethylase is controlled by the transcription factor CREB (cAMP-response binding protein), which is activated by cAMP/PKA-dependent phosphorylation ^70^. For instance, the beta2-adrenergic receptor agonist Clenbuterol engages cAMP/PKA/p-CREB signaling, which then drives the expression of *Jhdm2a*, a histone demethylase via direct binding of p-CREB to the CRE (cAMP response element) site in the JHDM2a promotor ^71^. For ADCY8, *de novo* expression seems to result from both, transcriptional activation and alternative splicing ^68^. Strikingly, the mouse *Adcy8* promoter contains a CRE site upstream of the TSS that is required for basal and cAMP-dependent promoter activity ^72, 73^. Thus, cAMP-dependent control of *Adcy* isoforms, directly or indirectly by epigenetic modifications, might have emerged as unifying mechanism.

Expression of truncated protein isoforms in metabolically active tissue, like adipose tissue or muscle, is conserved among different proteins that converge in common signaling pathways in a cell-type specific manner. Here, we demonstrated that cold exposure results in the expression of a truncated PPARGC1A-AT proteoform, which contributes to induction of *Adcy3-at* expression. The stimulus that induces *de novo* expression of the truncated proteins can, however, be different yet appropriate towards the seminal function of the tissue, i.e., cold exposure for the thermogenic BAT and endurance training for truncated PPARGC1A isoforms leading to the expression of truncated PPARGC1A isoforms PGC-1a2, 3, or 4^28, 61, 74^. As these stimuli, and the resulting protein expression, are crucial for the function and homeostasis of respective tissues, identifying pharmacological means to target these truncated isoforms is therapeutically relevant. Based on our results from *Adcy3ΔAT* mice, pharmacologically targeting *Adcy3-at* expression could pave the way to re-activate BAT function and thereby mitigate, or even prevent, obesity and obesity-related metabolic decline.

## Methods

### Animal care and research diets

Experimental animals were kept in individually ventilated cages (IVC Type II long) in a pathogen-free (SPF) animal facility with controlled temperature (22-24 °C), light/dark cycle (12h/12h) and humidity (50-70%). Care of animals was within institutional and animal-care committee guidelines approved by (1) local (Bezirksregierung Köln) or regional (Tierschutzkommission acc. §15 TSchG of Landesamt for Natur, Umwelt und Verbraucherschutz (LANUV) North-Rhine Westphalia, Germany) authorities, internal accession no. 84-02.04.2017.A009 or (2) Ministry of Environment of Denmark (Miljø- og Fødevarestyrelsen), internal accession no. 2018-15-0201-01562. All animals were maintained and regularly backcrossed to a C57BL/6N background and housed in groups of 3-4 animals per cage and had *ab libitum* access to food and drinking water. All mice were sacrificed by cervical dislocation or carbon dioxide asphyxiation. Unless otherwise indicated, animals were allowed ad libitum access to chow diet (ssniff® R/M-H Low-Phytoestrogen, V1554) containing 62 kJ% carbohydrates, 27 kJ% protein and 11 kJ% fat and drinking water. Diet-induced obesity (DIO) was achieved by feeding a high-fat diet (HFD, D12492 (I) mod; Sniff) containing 22 kJ% carbohydrates, 24k J% protein and 54 kJ% fat from starting at 6-8 weeks of age.

### Mouse husbandry

*Adcy3* floxed mice were kindly provided by Chen and colleagues ^24^. Herein, exon 3 of the *Adcy3* gene is flanked by two intronic *LoxP* sites, each 75 base pairs upstream or downstream of the exon. Deleting exon 3 of *Adcy3* causes a frame shift mutation, resulting in a premature stop codon within the *Adcy3* gene. Adipose tissue-specific deletion of both, *Adcy3-fl* and *Adcy3-at*, isoforms of *Adcy3* was achieved using cre recombinase-mediated excision of *LoxP*-flanked (‘floxed’) gene sequences. For this, mice floxed for *Adcy3* (*Adcy3^LoxP/LoxP^*) were interbred with mice expressing the *Adipoq*-cre recombinase under control of mature adipocyte specific *Adipoq* promote (*Adipoq-cre^+/^*cre) ^36^. *AdipoQ*-Cre mice were obtained from Jax (Stock no. #010803) and backcrossed to C57BL/6N for at least five generations. Resulting (*Adcy3^LoxP/LoxP^, Adipoq-cre^+/cre^* mice (*Adcy3*-*AdcKO*) were compared to (*Adcy3^LoxP/LoxP^, Adipoq-cre^+/+^*) littermates as controls (*LoxP*).

### Generation of Adcy3ΔAT mice

*Adcy3ΔAT* knock-out ES cells were generated using CRISPR/Cas9 technology using a modified pX330 plasmid vector ^78, 79^ containing two gRNAs and a GFP-puromycin selection cassette. The plasmid was kindly provided by Dr. Stefan Frank and was constructed at the Max-Planck Institute for Molecular Biomedicine (Münster, Germany). The knock-out strategy targeted 978 bp deletion of *Adcy3-at* starting exon (‘exon 2b’) and flanking regions from *Adcy3* genomic DNA. A CRISPR design online tool (http://crispr.mit.edu/) was used to generate *Adcy3-AT* specific single guide RNA (sgRNA) sequences. The list of sgRNAs and primers for generation of *Adcy3*ΔAT mice are given in Supplementary table S3. The quality score of the design pipeline represents the reliability of an on-target activity computed as 100%, minus a weighted sum of off-target hit-scores. A chosen gRNA with a high-quality score was cloned into the BbsI-digested pX330 vector, followed by the second gRNA, which was cloned using a SapI-mediated Golden Gate assembly method for the concatemerization of gRNA cassettes.

### Vector generation

For CRISPR/Cas9 construct generation, 1 μg of the pX330 vector was enzymatically digested with FastDigest BbsI (Thermo Fisher Scientific) for 30 min at 37 °C. The digested vector was purified using the MinElute PCR Purification kit (Qiagen) following the manufacturer’s instructions. gRNA was phosphorylated by a T4 polynucleotide kinase according to product instructions (New England Biolabs). For the amplification of phosphorylated sgRNA it was incubated with 1 μl of forward and reverse primers (100 μM, Supplementary Table S3) for 30 min at 37 °C. Annealing was performed at 95 °C for 5 min and subsequently the reaction was cooled to 25 °C at a cooling rate of 5 °C/min. 1 μl of the oligo reaction was ligated with 50 ng of digested pX330 vector and was assessed using the Quick Ligase kit (New England Biolabs) for 20 min at room temperature according to the manufacturer’s instructions. Plasmid Safe DNase kit (EpiCentre) was used to degrade any unspecific recombination products for 30 min at 37 °C, following the instructions of the kit.

### Culturing mouse embryonic fibroblasts and ES cells and ES cell transfection

Mouse embryonic fibroblasts (EF) were used as feeder cells for mouse embryonic stem cells (ESC). EF cells were passaged before reaching full confluence and EF-cell medium was changed to fresh medium every two days. To obtain mitotically inactive fibroblasts, cell growth was stopped by adding 10 μg/ml MMC after three generations (EF3) and the reaction was incubated for another 2-4 h. MMC was removed, and cells were rinsed with 1xPBS. The cells were afterwards stored by freezing or directly used for seeding. To seed MMC-treated EF3 cells, tissue culture dishes were gelatinized with 0.2% gelatin in advance to enhance the attachment of MMC-treated cells to the dish-surface. ES cells were cultivated on top of MMC-treated EF3 cells and ES cell culture medium was exchanged every day. ES cells were passaged to new culture plates or were frozen for long term use and to maintain pluripotency.

10^7^ ESC were mixed with 40 μg of the CRISPR construct dissolved in PBS, resuspended in 400 µL transfection buffer (RPMI without Phenol Red, Gibco, Karlsruhe, Germany), and mixed with the plasmid construct DNA. The total volume was adjusted to 800 μl. Transfection was achieved by electroporation at 450 μFD and 240 V at room temperature (Gene Pulser Xcell, Bio-Rad Laboratories GmbH, München, Germany) in a 4 mm electroporation cuvette. The time constant was set to around 7-10 seconds. After 5 min incubation, cells were resuspended in ES-cell medium, diluted 1:20000 and plated on a gelatinized 10 cm culture dishes with previously prepared MMC-treated EF3 cells. Following two days of incubation in ES-medium, single clones were transferred to three 96-well plates and selected via colony PCR. In addition to PCR confirmation, Southern blot analysis was used to confirm the deletion of *Adcy3-AT* as explained below. Validated colonies were transferred in 40 μl PBS to a 96-well (round-bottom) plate containing 25 μl trypsin. Trypsinization took place for 5 min at 37 °C and the reaction was stopped by adding 100 μl ES-medium. Cell suspension for each colony was added up to 15 mL with ESC-medium and seeded in one separate 10 cm dish per colony. To achieve higher homogeneity among targeted ESC populations, positive colonies were again seeded out after expansion in three 96-well plates each and selection by PCR and Southern blot repeated. Positive colonies were trypsinized and seeded on 10 cm dishes for expansion as before.

### Generation of Adcy3ΔAT chimeric mice from genetically modified ESC clones and genotyping

Positive *Adcy3ΔAT* clones were injected into donor blastocysts, which was performed by Taconic Biosciences (Cologne, Germany). Injected blastocysts were implanted into pseudo-pregnant foster mice and resulting progenies (Generation F1) partly develop from cells derived from the *Adcy3*ΔAT ESC. Male chimeric F_1_ mice were backcrossed to C57BL/6N mice to produce the F_2_ generation. Germline transmission was confirmed by PCR and Sanger sequencing of DNA extracted from the tails of F_2_ mice. Genotyping protocols for the *Adcy3ΔAT* transgenic allele was conducted using the following primers which were also used for genotyping from mouse tail DNA: *Adcy3ΔAT*_E2B_F_1: 5’-GGGACTGAGGGAGCCTAAGA-3’ *Adcy3ΔAT*_E2B_R: 5’-GGCCAGGTTACATGAGGACA-3’ *Adcy3ΔAT*_E2BF_2_R2: 5-’-GGAGAGCTTCGAGTGTGTCAAG-3’.

### Southern blot of Adcy3ΔAT embryonic stem cells

For confirmation of *Adcy3-at* deletion in ESC, genomic DNA (5-10 μg) derived from transfected ESC was digested by high concentrated HindIII-HF or high concentrated BamHI-HF (NEB). In all Southern blot analyses, wildtype Bruce 4 (Br4) ESC DNA was used as negative control. Overnight digested DNA was loaded with loading dye onto a 0.8 % agarose gel and separated at 30 V for at least 16 hours. The DNA was consequently de-purinated within the gel by incubating the gel in 0.25 M HCl for 20 min under continuous shaking. DNA was then transferred into a charged Amersham Hybond-XL nylon membrane by alkaline capillary transfer using a sodium hydroxide solution (0.4 M NaOH) and the membrane was incubated in 2X SSC for 20 min and finally dried at 80 °C for 45 min to fix the DNA on the membrane. To probe for the modified *Adcy3ΔAT* allele, a 394 bp probe was amplified from Bruce4 ESC genomic DNA with primers 5’-ATCCTGTGCTGACATGGGTG-3’ and 5’-AATTCCCCACTGACCAACGG-3’ using high-fidelity PCR Master Kit. The extracted probes were radioactively labeled with 2.5 μl [alpha-32P] dCTP (Amersham) according to the instructions of the kit (LaddermanTM Labeling Kit, Takara). The probe was mixed with 100 μl TE buffer (10mM, pH8.5) and purified via spin-column chromatography using Illustra MicroSpin S-200 HR columns. The dried membrane was incubated in 2X SSC buffer to prepare for pre-hybridization. Pre-hybridization was performed in pre-hybridization buffer (1 M sodium chloride, 50 mM Tris-CL pH 7.5, 10 % Dextran sulfate, 1% SDS, 250 μg/ml salmon sperm DNA) at 65 °C for at least 4 h. After pre-hybridization radioactively labeled probes were added to the membrane and the reaction was incubated overnight at 65 °C. After several washing steps, the membrane was placed in direct contact with Kodak MS hypersensitive films for 4-24 h to detect the radioactive signal. Expected product sizes were 4.75 kb for *Adcy3* wildtype and 3.75 kb for *Adcy3ΔAT* constructs.

### Plasmid vector cloning

Overexpression constructs (ADCY3-AT, 3201 bp and ADCY3-FL, 3792 bp) were PCR amplified from synthetic oligonucleotides by Thermo Fischer scientific. The DNA fragment was inserted into pcDNA3.1(+) vector backbones using NheI / XhoI restriction enzymes. For ADCY3-AT the first ATG in-frame within exon 2 was used as start codon. Sequences encoding Kozak sequences (GCCACC) and a 295 bp artificial intron sequence were included upstream of ATG. To generate pc3.1(+)-mADCY3-AT-FLAG, the FLAG tag was added by PCR and the coding sequence cloned into the vector backbone using EcoRI/XhoI. Additional details on cloning procedures are available upon reasonable request.

### Illumina and Nanopore RNA sequencing and data analysis

Paired end libraries were constructed using the NEBNext Ultra II RNA Library Prep Kit (New England Biolabs, Ipswich, MA, USA) following the manufacturer’s protocol and sequenced on a NovaSeq 6000 (Illumina, San Diego, CA, USA) in 2×50-bp paired-end reads. Transcriptional expression was quantified using Salmon^80^ by mapping RNA-seq reads to the ENSEMBL m38 (mm10) transcriptome. Adcy3-at transcript identified in the de novo ONT/Illumina transcriptome assembly (see section below), was added to the ENESEML m38 transcriptome and counted as an independent gene. Transcript counts were imported and summarized to ENSEMBL genes IDs using tximport^81^ and differential expression analyses were conducted with edgeR ^82^using glmQLFTest. KEGG pathway enrichment analyses were conducted using clusterProfiler package for R^83^.

### Nanopore RNA-Sequencing and data processing

RNA was isolated by phenol chloroform extraction and alcohol precipitation followed by two consecutive rounds of poly-A selection using oligo(dT) beads (GenElute mRNA Miniprep Kit, Sigma MRN10) following the manufacturer’s recommendations. Subsequently, RNA was alcohol precipitated using NaAcetate and glycogen following the protocol from the Ribo-Zero rRNA Removal Kit (Illumina). Nanopore sequencing libraries were prepared using the TeloPrime Full-Length cDNA Amplification Kit (Lexogen, Austria), the SQK-DCS109 direct cDNA sequencing kit (Oxford nanopore technologies, ONT) and the SQK-RNA002 direct RNA sequencing kit (ONT) according to manufacturer’s protocols. Sequencing was performed using FLO-MIN106 R9 flow cells on the GridION platform (ONT). Reads were mapped against the murine genome (GRCm38.p6) using minimap2 ^84^ and transcriptomes reassembled using FLAIR ^75^. Tracks were visualized using gviz ^85^.

### Single-nucleus RNA-Sequencing (snRNA-seq) data processing

Pre-processed data from snRNA-seq from 10X sequencing of mouse 1°BA housed at room-temperature, cold-exposure and/or thermoneutrality was adopted from Sun^34^ (ArrayExpress using accession code E-MTAB-8562. Data was normalized (SCTransform), analyzed and plotted using Seurat 4.0.1 package for R^80–83, 86^.

### Chromatin Immunoprecipitation (ChIP) coupled to sequencing (ChIP-seq) and ChIP-qPCR

Prior to ChIP-seq, brown adipose tissue was dissociated using a gentleMACS Dissociator (Miltenyi Biotec, Germany). The cell suspension was crosslinked with 1 % formaldehyde for 10 min at RT and the reaction was quenched with 0.125 M glycine for 5-10 min at RT. Cells were washed 2X with cold PBS and PMSF and snap-frozen in liquid nitrogen before storing at −80°C. For histone posttranslational modification (PTM) sequencing, BAT of two mice were used and steps performed on ice. First, frozen pellets were thawed on ice for 30-60 min. Pellets were resuspended in 5 ml LB1 (50 mM Hepes, 140 mM NaCl, 1 mM EDTA, 10 % Glycerol, 0.5 % NP-40, 0.25 % Triton X-100) by pipetting and then rotated vertically at 4°C for 10 min. Pellets were centrifuged for 5 min at 1350 x g at 4°C (GH-3.8 rotor = 2400 rpm) and supernatant was aspirated. Pellets were resuspended in 5 ml LB2 (10 mM Tris, 200 mM NaCl, 1 mM EDTA, 0.5 mM EGTA) and incubated at vertical rotation and at room temperature for 10 min. Samples were centrifuged for 5 min at 1350 x g at 4°C and supernatant was carefully aspirated. Then, samples were resuspended in 3 ml LB3 (10 mM Tris, 100 mM NaCl, 1 mM EDTA, 0.5 mM EGTA, 0.1 % Na-Deoxycholate, 0.5 % N-lauroylsarcosine) and were separated into 2 x 1.5 ml in 15 ml polypropylene tubes, in which they were sonicated with the following settings by Bioruptor Plus sonication: Power = high, ‘on’ interval = 30 sec, ‘off’ interval = 45 sec, total time = 10 min (18 cycles of ‘on’/’off’). Sonicated samples were transferred to a 1.5 ml microfuge tube and were centrifuged for 10 min at 16000 x g at 4°C to pellet cellular debris. 10 % of sample solution were stored to be used as input control, while the rest was used for ChIP. To capture different H3K4me3 modifications, 3 µl of anti-H3K4me3 antibodies (Active Motif, cat. No 39159) were added to the sonicated ChIP reaction and rotated vertically at 4° C overnight. The next day, 100 μl Dynabeads (Protein A or Protein G) for each ChIP sample were prepared according to the manufacturer’s instructions, mixed with 1 ml of antibody-bound chromatin, and rotated vertically at 4°C for at least 2-4 hours. Bound beads were washed at least five times in 1 ml cold RIPA and once in 1 ml cold TE buffer containing 50 mM NaCl. Samples were eluted for 15 min with elution buffer at 65°C and continuously shaken at 700 rpm. Beads were separated using a magnet and 200 μl supernatant were transferred to fresh microfuge tubes. Previously stored input samples were thawed and mixed with 300 μl Elution buffer. ChIP/input samples were incubated at 65°C in a water bath overnight to reverse the crosslinking reaction. On the third day, 1 volume TE buffer was added at room temperature to dilute SDS in both ChIP and input samples. For digestion of RNA and protein contamination, RNase A was added to the samples and incubated in a 37°C water bath for 2 hours; then proteinase K was added to a 0.2 mg/ml final concentration and incubated in a 55°C water bath for 2 hours. Finally, DNA was extracted using a standard phenol-chloroform extraction method at room temperature and DNA concentrations were measured using a NanoDropTM ND-1000 spectrophotometer or QubitTM dsDNA HS Assay Kit and stored at −80 °C until sequencing or ChIP-qPCR.

H3K4me3-immunoprecipitated genomic DNA was diluted 1:100 and Chip-qPCR performed with a Light Cycler 480II machine (Roche) using technical duplicates and ChIP-qPCR signals were calculated as % of input. Standard deviations were calculated from technical duplicate reactions and represented as error bars. Quantitative PCR primer sequences were:

*Adcy3-fl* TSS_F: 5’-GATGGACTTCCACGAGGCTG-3’

*Adcy3-fl* TSS_R: 5’-TACTTCCTTTCCCCCACCCA-3’

*Adcy3-at* TSS_F: 5’-GGAGAGCTTCGAGTGTGTCAAG-3’

*Adcy3-at* TSS_R: 5’-GTCTCACCTTAAGGCTCCTCCT-3’

*Adcy3 exon 3*_F: 5’-CTGTGCCAGATTGTCTCCGT-3’

*Adcy3 exon 3*_R: 5’-GTCATGGACTTGGGCTTCCA-3’

### Mouse tissue RNA isolation

RNA from indicated tissues and primary brown and inguinal white adipocytes was isolated using Trizol® according to manufacturer’s protocols for total RNA isolation.

### CHO K1 cell culture

CHO K1 WT cells (ATCC, #CCL-61) were maintained in F-12 Nut Mix + GlutaMAX (#31765-027, Thermo Fisher) at a subconfluent level and passaged every 4 days. Cells stably expressing ADCY3-FL-HA were kept in medium containing 0.8 mg/ml Geneticin (G418 Sulfate, Gibco).

### Isolation of depot-specific stromal-vascular fraction (SVF) derived 1° adipocytes

Inguinal white and intrascapular brown adipose tissues from 6-8 weeks-old male mice were dissected after carbon dioxide asphyxiation, minced, and digested with collagenase II (2 mg/mL), DNAse I (15 kU/mL), Dispase II (for BAT: 1.5 mg/mL) and 3.4 % BSA supplemented serum-free SVF culture media Dulbecco’s Modified Eagle Medium:Nutrient Mixture F-12 (DMEM/F-12) pre-supplemented with 1 % P/S, 0.1 % Biotin and 0.1 % pantothenic acid. Minced tissues incubated in 37 °C shaker at 120 rpm/min for 45 minutes and pipetted in every 15 minutes until the minced tissue particles were completely dissolved. SVF fraction cells were enriched in the pellet of the tubes by sequential filtering and washing steps with 250 μm and 70 μm sterile cell strainers in 10 % FBS supplemented SVF culture media. Enriched SVF cells seeded in 6-well plate wells (1 well/mice) with 20% FBS supplemented SVF culture media and incubated in cell culture incubator for the required amount of time to enable expansion of the culture.

### Induction of SVF adipogenesis

iWAT and BAT depot-specific stromal vascular fraction (SVF) at 80 % confluency were induced for adipogenic differentiation with induction cocktail (1 μM Rosiglitazone, 850 nM Insulin, 1 μM Dexamethasone, 250 μM IBMX, and, for only BAT, 125 μM Indomethacin and 1 nM T3) in 10 % FBS supplemented SVF culture media. After incubation for 2 days with induction media, culture media replenished with the differentiation media (1 μM Rosiglitazone, 850 nM Insulin and for only BAT with 1 nM T3 in 10 % FBS supplemented culture media) on days 2, 4, and 6. Cells were treated with compounds (10 μM CL-316,243; 2 μM Forskolin) on day 7 for and harvested the same day after compound stimulations.

### LNA-mediated gene knockdown in primary adipocytes

For transfection of LNA GapmeRs in 1°BA and 1°iWA, 80,000 cells were seeded per well in 6-well plates and grown to confluence in growth medium. Immediately before transfection, medium was changed to fresh differentiation medium without antibiotics. In the meantime, antisense LNA GapmeRs (Exiqon A/S) were diluted to the concentration of 100 nM in OptiMEM and transfected using Lipofectamine® 2000 according to the manufacturer’s instructions. LNA sequences are provided in Supplementary Table S5.

### AAV-mediated cre transduction in primary adipocytes

On day 4 of iWA and BA differentiation, committed adipocytes were trypsinized, counted and seeded into multi well plates with the experimental setup (180.000 cells / cm^2^ growth area) with the differentiation media containing AAV viruses. Primary adipocytes were transduced with AAV8-CMV-Cre (pENN.AAV.CMVs.Pl.Cre.rBG, Addgene #105537-AAV8 Cre) and AAV8-CMV-eGFP (pAAV.CMV.PI.EGFP.WPRE.bGH, Addgene #105530-AAV8 eGFP) viruses with the amount of 100. 000 MOI. Virus containing media replenished on day 6 with fresh differentiation media and cells were treated with compounds (10 μM CL-316,243) on day 7 for and harvested the same day after compound stimulations.

### Determination of oxygen consumption rates and measurement of glycolytic activity

*Adcy3-at* deficient SVFs from indicated adipose depots of transgenic mice were seeded into Agilent Seahorse XFe96 Bioanalyzer microplates. Per well, 50,000 cells were seeded and incubated in DMEM/Ham’s F-12 medium plus 10 % Fetal Calf Serum, 1 % P/S, 0.1 % Biotin and 0.1 % Pantothenic acid (Growth medium) at 37° C and 5 % CO2 at a standard incubator until confluency is reached. To induce commitment of SVFs into mature adipocytes within Xfe96 microplates, freshly prepared 0.05 % Insulin, 0.005 % Dexamethasone, 0.001 % Rosiglitazone and 0.05 % IBMX (1°iWA) or 0.1 % Indomethacin, 0.001 % Triiodothyronine (1°BA) in growth medium (induction medium) were added. After 48 h of induction, differentiation was initiated using freshly prepared 0.001 % Rosiglitazone (1°iWA) or 0.001 % Triiodothyronine (1°BA) in growth medium (differentiation medium). Differentiation was achieved after 3-4 days of incubation in the differentiation medium. For each seahorse plate the corresponding calibration plate was prepared 24 h prior to experiments using 200 µl XF Seahorse Calibrant Agilent per well. The plate was incubated for 24 h in a non-CO2 incubator at 37° C and the instrument set to 37° C 24 h prior to the experiment. One hour before measurements, plates were washed with 1xPBS and the medium changed according to the corresponding experiment analyte kits (MitoStressKit or GlycoStressKit, provided by the manufacturer). Prior to measurement, calibration was started using calibration plates, measuring O2 and pH LED Value/emission/Initial reference Delta for each well. After calibration cartridges were kept within the machine and measurement of adipocyte-containing microplates commenced. Measurement parameters were mix 3 min, wait 0 min, measure 3 min with each reagent’s effect assessed within three (MitoStressKit) or four consecutive measurement cycles (GlycoStressKit) with a total duration of 18 min or 24 min per reagent injection. All measurements started with measuring basal values, followed by injection of Oligomycin, FCCP and Rotenone plus Antimycin A (MitoStressKit) or Glucose, Oligomycin and 2-Deoxy-Glucose, GlycoStressKit). (1) For MitoStressKit, corresponding media were prepared before the experiment and consisted of Basal Seahorse Medium supplemented with 25 mM Glucose, 1 mM Glutamine, 2 mM Sodium Pyruvate, set to pH = 7.4 and filtered sterile. Per Plate ca. 25 ml of MitoStress Medium were needed and Seahorse cell plates were changed to 180 µl MitoStress medium 1 h prior to calibration in a non-CO2 incubator at 37° C. The calibration plate possessed a cartridge having 4 pockets per well. Before the measurement pocket A was filled with 20 µl 10 µM Oligomycin, pocket B with 22 µl 10 µM FCCP and pocket C with 25 µl 5 µM Antimycin A and Rotenone. (2) For GlycoStressKit, the cell plate was washed with 1xPBS, and media changed to filtered 180 µl GlycoStressKit medium. GlycoStressKit medium consisted of Basal Seahorse Medium supplemented with 1 mM Glutamine and 2 mM Sodium Pyruvate, set to pH = 7.4 and stored for 1 h in a non-CO2 incubator at 37° C. The calibration plate possessed a cartridge having 4 pockets per well. Shortly before the measurement pocket A was filled with 20 µl 10 mM Glucose, pocket B with 22 µl 10 µM Oligomycin and pocket C with 25 µl 50 mM 2-Deoxy-Glucose.

### Intraperitoneal glucose tolerance test (GTT) and insulin tolerance test (ITT)

GTT was carried out at 12am after a 6 h fast starting in the morning. After determining basal blood glucose levels (0min), animals received an intraperitoneal bolus of 2 g glucose per kilogram of body weight (20 % glucose, Delta select). Blood glucose levels were determined 15, 30, 60 and 120 min after injection using an automatic glucose monitor (Contour, Bayer Diabetes Care). ITT was carried out in random-fed mice at 9-10am in the morning in fresh cages with bedding, free access to drinking water but no food. After determining basal blood glucose levels (0min), each animal received 0.75 U/kg of body weight of insulin (Actrapid; Novo Nordisk). Blood glucose levels were recorded after 15, 30, 60 and 120 min. Those animals that showed no increase/decrease of blood glucose levels after i.p. injection of glucose or insulin, assuming injection outside of the peritoneal cavity as required for the assay, were excluded from analysis.

### Indirect calorimetry (PhenoMaster)

Indirect calorimetry (PhenoMaster, TSE Systems GmbH, Bad Homburg, Germany) was used to evaluate energy expenditure (EE), locomotor activity (LMA), respiratory exchange ratios (RER) as well as food and water intake (1) in the animal facility of Max Planck Institute for Metabolism Research or (2) the Biomedical Laboratories (University Hospital Odense, OUH). Mice were allowed to acclimatize to single housing conditions in metabolic cages for two to four days. Food and water were provided *ad libitum* during acclimatization and throughout all experimental measurements. All parameters of indirect calorimetry were measured for a period of 1-10 days depending on the duration of cold exposure. Oxygen consumption and carbon dioxide production were measured in units of volume every 10 min for indicated amounts of days (including measurements for both cold and warm ambient temperatures) to determine the respiratory quotient (RQ = VCO2/VO2 2/VO2))) × 4.1868). LMA was measured by a multidimensional infrared light beam system. Integrated scales assessed water and food intake at 2 min intervals and automatically calculated cumulative water and food intake. During measurements in the PhenoMaster System, mice were initially kept at a constant temperature of 23 °C for 72 h, followed by 4 °C for 1-6 days depending on the experimental setup.

### mRNA isolation and quantitative RT-PCR (qPCR) analysis

mRNA isolation from primary adipocytes and adipose tissues were lysed within a monophasic solution of phenol and guanidine isothiocyanate reagent (TRI Reagent®, Sigma-Aldrich #T9424) and eluted with RNAse/DNAse free water following silica-based RNA spin column enrichment. RNA concentrations measured by spectrophotometer and 500 ng RNA used for reverse transcription (High-Capacity cDNA Reverse Transcription Kit, Thermo Fisher #4368813). Abundance of *Adcy3-at*, *Adcy3-fl*, *Adipoq*, *Cidea*, *Dio2*, *Elovl3*, *Emr1*, *Fabp4*, *Lsgals3*, *Pparg* isoform 2 (*Pparg2*), *Ppargc1a-at*, *Ppargc1a-fl*, *Prdm16* and *Ucp1* were quantified using SYBR Green based quantification method (FastStart Universal SYBR Green Mastermix, Roche® Life Science #4913914001) and mRNA abundance was calculated using relative quantification methods (2^−ΔΔCT^). Transcript levels of mRNAs were normalized to hypoxanthine phosphoribosyltransferase 1 (*Hprt1*) or general transcription factor IIB (*Gtf2b*) expressions. Primer sequences for SYBR Green based quantification are provided in Supplementary Table 4.

### Protein isolation

To isolate total protein from 1°BA and 1°iWA, 100-250 μl RIPA buffer were added to frozen cells in cell culture plates. Using a cell scraper, the adipocytes were detached and transferred into 1.5 ml reaction tubes. In case of adipose tissue, Precellys® zirconium oxide beads and 500-1000 μl RIPA buffer was used to disrupt and homogenize adipose tissue using a FastPrep-24TM 5G Homogenizer (mouse muscle program: 3x 90 sec). To disrupt the cell membrane of the tissue and primary cells, tubes were snap frozen in liquid nitrogen and thawed on ice three times. The samples were centrifuged for 10 min at 12000 x g and at 4 °C. The supernatant was transferred to a new reaction tube and samples were stored at −80 °C. Quick StartTM Bradford Protein Assay Kit and Pierce BCA Protein Assay Kit were used for the quantification of protein concentrations according to the instructions of the kits. A FilterMaxTM F5 microplate reader was used to measure protein concentrations at 595 nM.

### SDS-PAGE and Western Blot analysis

Protein samples were diluted in 4X Laemmli Buffer (Bio-Rad Laboratories) containing 5 % ß-Mercaptoethanol and were boiled at 96 °C for 6 min (unless when probing for ADCY3). Protein lysates were separated based on their molecular weight by sodium dodecyl sulfate-polyacrylamide gel electrophoresis (SDS-PAGE). Samples and PageRulerTM Prestained Protein Ladder (ThermoScientific) were loaded onto Mini-PROTEAN® precast gradient (4-20 %) polyacrylamide gels (Bio-Rad Laboratories) and proteins were separated in an electric field of 90-120 V for varying time spans in SDS Running Buffer (25 mM Tris-base, 0.192 M Glycine, 0.1 % SDS).

SDS-PAGE separated proteins were transferred to methanol-activated PVDF membrane by wet transfer. Wet transfer was performed in a blotting chamber with ice-cold transfer buffer (48 mM Tris-base, 39 mM glycine, 0.037 % SDS and 20 % methanol) for 2-3 h at 120 mA per membrane. After protein transfer membranes were incubated in Ponceau solution (Sigma-Aldrich) to confirm even transfer. Afterwards, membranes were rinsed once in TBS-T (500 mM Tris base, 1.5 M NaCl, 1 % Tween-20, pH=8) and incubated for 1 h in a Western Blocking Reagent solution (Sigma-Aldrich). Membranes were incubated overnight at 4 °C with 20 ml of the respective primary antibody solution. Primary antibodies were anti-HSC70 (sc-7298, Santa Cruz Biotechnology, dilution 1:10,000), anti-UCP1 (#14670, Cell Signaling Technology, dilution 1:1000), anti-ADCY3 (#Ab14778, Abcam, dilution 1:500, and anti-phospho-PKA Substrate (#9624, Cell Signaling Technology, 1:1000). Membranes were washed three times for 10 min with TBS-T and incubated for 1 h at room temperature with 10 mL of the respective horseradish peroxidase (HRP)-coupled secondary antibody. Wash steps were repeated, and 1 mL of chemiluminescent substrate (#1705061, Biorad) was applied on the membrane and visualized by Amersham Imager 680.

For CHO cells, 4x sample buffer (200 mM TRIS/HCl pH 6.8, 8 % (w/v) SDS, 4 % (v/v) β-Mercaptoethanol, 50 % (v/v) glycerin, 0.04 % (w/v) bromphenolblue) was added to the samples at a final concentration of 1x, directly loaded on an 8.75 % SDS-PAGE gel, and separated by molecular weight. Subsequently, proteins were transferred onto a methanol activated PVDF membrane by semi-dry transfer. The membrane was blocked using PBS Intercept blocking buffer (LI-COR) for 30 min at room temperature. The membrane was then incubated with the primary antibody solution overnight at 4°C. The following primary antibodies were used: anti-FLAG (1:2000, F1804, Sigma-Aldrich) and anti-HA (1:1000, Roche, clone 3F10). Following primary antibody incubation, the membrane was washed 3 times with PBS-T for 10 min at room temperature. The membrane was then incubated with the respective IRDye secondary antibodies (1:20000, LICOR) for 1 h at room temperature. After washing the membrane 3 times with PBS-T and once with PBS, protein bands were imaged using a LI-COR imaging system. Protein bands were quantified using ImageJ.

### Co-immunoprecipitation

2.1 x10^6^ CHO K1 cells stably expressing ADCY3-FL-HA were seeded on 10 cm cell culture dishes (Greiner). The following day, the cells were either transfected with pc3.1-ADCY3-AT-FLAG or pc3.1Zeo-mCherry using polyethylenimine (Sigma-Aldrich). To increase expression of the transfected constructs, 5 h after the transfection sodium butyrate was added to the cells to a final concentration of 5 µM. 24 h later, cells were washed with 10 mL PBS and scraped in 1 mL PBS. The cell solution was then transferred into a precooled tube and centrifuged at 500 x g for 5 min at 4°C. The supernatant was aspirated, and the cell pellet resuspended in 300 µL lysis buffer (20 mM TRIS/HCl pH 8, 137 mM NaCl, 2 mM EDTA, 1% NP-40, 1:500 mammalian protease inhibitor cocktail (Sigma Aldrich)). Following 60 min incubation on ice, the samples were centrifuged at 10000 x g for 10 min at 4°C. The supernatant was transferred into a new tube and the protein concentration was determined using the Pierce BCA Protein Assay Kit (Thermo Scientific). Protein concentration was adjusted to 0.5 µg/µL using lysis buffer and 45 µL were stored at −80°C (“input”). To reduce the amount of unspecific binding, the lysate was incubated with uncoupled NHS-activated magnetic beads (Thermo Fisher). 40 µL NHS-activated beads were equilibrated in 400 µL equilibration buffer (10 mM TRIS/HCl pH 7.4, 150 mM NaCl) and separated from the supernatant using a magnet. The beads were resuspended in 500 µL of the 0.5 µg/µL lysate and incubated for 1 h at 4°C end over end. Beads and lysate were then magnetically separated, and 45 µL of the supernatant were stored at −80°C (‘non-bound 1’). Magnetic anti-FLAG-M2 beads (Sigma-Aldrich) were equilibrated in 400 µL equilibration buffer (10 mM Tris/HCl pH=7.4, 150 mM NaCl), resuspended in the supernatant, and incubated end-over-end overnight at 4°C. The next day, beads and supernatant were magnetically separated, 45 µL of the supernatant were stored at −80°C (‘non-bound 2’), and the rest of supernatants were discarded. The beads were washed 4 times with 375 µL wash buffer (20 mM TRIS/HCl pH8, 137 mM NaCl, 2 mM EDTA, 0.2 % NP-40, 1.500 mammalian protease inhibitor cocktail). 45 µL of wash fractions 1 and 4 were saved for Western blot analysis. Proteins bound to the beads were eluted by incubating the beads in 75 µL elution buffer (0.1 M glycine pH 3.0) for 3 min at room temperature. Subsequently, 20 µL 1 M TRIS/HCl pH 8 were added for neutralization. 45 µL of the neutralized elution fraction were analyzed by western blot analysis.

### Biotinylation assay

The biotinylation assay was performed using the Pierce Cell Surface Biotinylation and Isolation kit (A44390, Thermo Scientific). Cells were treated with the biotinylation reagent Sulfo-NHS-SS-Biotin according to the manufacturer’s protocol, harvested, and lysed. The lysate was transferred to a column containing a NeutrAvidin-agarose-resin and incubated end-over-end overnight. The column was centrifuged and the flow-through (sample F) was collected. The column was washed three times (samples W1, W2, W3) with 500 µL wash buffer and the labeled protein was eluted (sample E) using 200 µL 4x SDS sample buffer containing 50 mM dithiothreitol (DTT). After elution, beads of the agarose matrix were scraped off the column, boiled in 100 µL 4x SDS probe buffer and centrifuged. For each fraction, 40 µL were loaded.

### Membrane sheets and immunofluorescent labeling

CHO cells were seeded on poly-L-lysine (PLL, 0.1 mg/ml, Sigma Aldrich, #P1399-100MG)-coated 13 mm glass coverslips (VWR) coated glass coverslips in a 4-well dish (VWR) and cultured for 24 h. Before sonification, cells were washed with PBS and sonicated in 500 µl sonification buffer (120 mM glutamate, 20 mM potassium acetate, 10 mM HEPES, 10 mM EGTA, pH 7.2 using the VibraCell Sonifier (0.1 s pulse, distance to cover slip: 3 mm, amplitude: 1-20 %). Cells were fixed with 4 % paraformaldehyde (Alfa Aesar, ThermoFisher Scientific, #43368) for 10 min at room temperature. After washing thrice with PBS, cells were blocked with CT (0.5% Triton X-100 (Sigma Aldrich, #X100) and 5% ChemiBLOCKER (Merck Millipore, #2170) in 0.1 M NaP, pH 7.0) for 30 min at room temperature. Primary and secondary antibodies were diluted in CT and incubated for 60 min each at room temperature, respectively. Coverslips were mounted with one drop of Aqua-Poly/Mount (Tebu-Bio, #07918606-20). The following primary antibodies were used: anti-calnexin (rb, 1:200, Sigma Aldrich, #C4731), anti-HA (rt, 1:1000, Roche, clone 3F10), anti-CNG channel (ms, 1:100, clone 3B10) (Pichlo et al, JCB 2014). As a DNA counterstain, cells were labeled with DAPI together with the secondary antibody (4’,6-Diamidino-2-Phenylindole, Dihydrochloride, 1:10.000, Thermo Fisher Scientific, #D1306). The following secondary antibodies were used: goat-anti-rabbit-Alexa647 (1:500, Thermo Fisher Scientific, #A21245), goat-anti-mouse-Alexa594 (1:500, Thermo Fisher Scientific, #A21125), goat-anti-rat-Alexa488 (1:500, Thermo Fisher Scientific, #A11006).

### AlphaFold modeling of Adcy3 structures

Folding prediction of ADCY3-FL and ADCY3-AT protein monomers and dimers were carried out with LocalColabFold based on AlphaFold2 using standard parameters ^54, 87^. ATP was docked to ADCY3 models by aligning catalytic domains with B chain of x-ray crystal structure PDB ID 3C16 (Catalytic Domain of Mammalian Adenylyl Cyclase). Computation of the models was performed on the UCloud interactive HPC system, which is managed by the eScience Center at the University of Southern Denmark. Molecular graphics performed with UCSF ChimeraX^88^. Alignment of structures was performed using ‘*mmaker*’ function in ChimeraX.

### PamGene serine-threonine kinase (STK) kinome arrays

Frozen brown adipose tissue was homogenized in 150 µL cold M-Per (Thermo Fisher) containing 1:50 HALT Protease and Phosphatase Inhibitor Cocktail (EDTA-free, 100x, Thermo Fisher) using a small, precooled pistil. Following 10 min incubation on ice, samples were centrifuged at 10000 x g for 5 min at 4°C. The liquid phase was carefully transferred into a new precooled tube, and this process was repeated at least twice (or until no lipid phase was left). Subsequently, the cleared lysates were aliquoted in 15 µL aliquots, frozen in liquid nitrogen, and stored at −80°C. Not more than 4 samples were handled simultaneously to avoid prolonged handling times. After the protein concentration was determined using Pierce BCA Protein Assay Kit (Thermo Scientific), 10 µg of protein was used as input and the STK assay was performed according to the standard protocol provided by PamGene. Kinome trees have been generated using the following software: http://phanstiel-lab.med.unc.edu/CORAL/

### Catch point cAMP assay (ELISA)

Brown adipose tissue cAMP levels were measured using the CatchPoint cAMP Fluorescent Assay Kit (Molecular Devices). A small piece of frozen BAT (2 – 5 mg) was transferred into a BeadBug tube (Sigma Aldrich) containing 1.0 mm zirconium beads (Merck) and 300 µL cold catchpoint lysis buffer. The tissue was disrupted using a BeadBug microtube homogenizer (Sigma Aldrich) at 400 rpm for 30s. Subsequently, samples were placed on ice for 1 min and homogenization was repeated twice. Following 15 min incubation at 4°C, each sample was sonicated 3 times for 1 min with 1 min incubation on ice in between each cycle. To clear the lysate from debris and insoluble lipids, samples were centrifuged at 10000 x g for 10 min at 4°C. The liquid fraction of the lysate was then transferred into a new tube and the process was repeated twice. The protein concentration of the cleared lysate was determined using the Pierce BCA Protein Assay Kit (Thermo Scientific). The CatchPoint cAMP ELISA was then performed according to the manual using 0.8 µg protein as input. Colorimetric measurement was performed after 30 min final incubation using a FLUOstar Omega (BMG Labtech).

### Human primary brown and white adipocytes and study characteristics

Deep neck BAT biopsies were acquired from a 52-year-old, non-diabetic female donor (BMI 24.1) undergoing thyroid surgery after giving written informed consent and approval by the ethics commitee of the University Hospital Bonn (Vote 076/18). Primary brown adipocytes were isolated according to protocols described recently (Jespersen et al., 2013) and cultured in 60mm culture dishes containing DMEM/F12, 10% FBS, 1% Penicillin/Streptomycin (all from Invitrogen) and 1nM acidic FGF-1 (ImmunoTools). Cells were incubated at 37°C with 5% CO2. Adipocytes were induced two days after full confluence with DMEM/F12 containing 1% Penicillin/Streptomycin, 0.1 mM dexamethasone (Sigma-Aldrich), 100 nM insulin, 200 nM rosiglitazone (Sigma-Aldrich), 540 mM isobutylmethylxanthine (Sigma-Aldrich), 2 nM T3 (Sigma-Aldrich) and 10 mg/mL transferrin (Sigma-Aldrich). After three days of differentiation, isobutylmethylxanthine was removed from the cell culture media. The cell cultures were left to differentiate for an additional nine days. Cells were treated with and without 1 µM NE for 16h. Total RNA was isolated using Trizol (Invitrogen). Reverse transcription was performed using ProtoScript II (NEB). qPCR reactions were assembled with Luna Master Mix (NEB) and qPCR was performed using a HT7900 (Applied Biosystems). Expression levels were calculated as delta Ct values relative to house-keeping gene hTBP (human TATA-box binding protein) serving as control.

### Comparative sequence conservation analyses of Adcy3-at and Ppargc1a-at genomic sequences

Analyses of alternative transcription (AT) start sites sequence conservation were conducted by first acquiring genomic contigs spanning either *Ppargc1a* or *Adcy3* via NCBI BLASTs using mouse or human transcript variants (accession numbers: NM_001377131.1; NM_138305.3; NR_132764.1) as query against whole genome shotgun contigs from representative species. Accession numbers of acquired contigs are listed in Supplementary Table S2. Contigs were imported into Geneious software version 9.1.8 (Dotmatics), and conserved exons were annotated using the ‘transfer annotations’ function. Alternative start sites from the mouse or human were then used to search for homologous sequences among contigs of various species using either the ‘transfer annotations’ function or EMBOSS 6.5.7 dotmatcher dot plots. MUSCLE alignments of conserved putative AT regions were assembled in Geneious and pairwise percent nucleotide identities were calculated relative to AT sites of either the mouse or human. These conservation levels were then mapped to the respective phylogenetic trees using the colour gradient function on the Interactive Tree of Life webserver^89^. The phylogenetic relationships and branch lengths illustrating the conservation of *Ppargc1a-at* were based on ^77^, while phylogenetic relationships and branch lengths for *Adcy3-at* conservation analyses were based on ^90^ for primates and ^91^ for rodents.

### Data availability

NGS datasets are currently being uploaded to Gene Expression Omnibus (GEO) and are provided upon reviewer’s request.

### Public datasets

H3K4me3 ChIP-seq from BAT of chow diet and HFD fed, male C57BL/6N mice housed at 22°C or exposed to 5°C for 24 h were downloaded from GEO (GSE20065). snRNA-seq data from *Adipoq*-tdTomato-positive adipocyte nuclei were downloaded from Array Express (E-MTAB-8562).

## Acknowledgements

We thank Jens Alber and Ann-Brit Marcher for indirect calorimetry (TSE Phenomaster) support. Sequencing was carried out at the Functional Genomics and Metabolism Research Unit, University of Southern Denmark. The authors thank Tenna P. Mortensen, Maibrith Wishoff, and Ronni Nielsen for sequencing assistance. We thank Catharina Baitzel and Anke Lietzau, Max-Planck Institute for Metabolism Research, for assistance in generating *Adcy3ΔAT* mice. JWK and BDL received funding from the University of Southern Denmark and the Danish Diabetes Academy, which is funded by the Novo Nordisk Foundation. JWK, AJ, and ARAAT received support from Challenge (#33444) and Bioscience and Basic Biomedicine Programs of the NNF (#28416). HT is supported by a postdoctoral fellowship form the DDA. SK was supported by a German Academic Exchange Service (DAAD) PhD scholarship (A/12/97620). We to thank the Microscopy Core Facility of the Medical Faculty at the University of Bonn for providing help, services, and devices funded by the Deutsche Forschungsgemeinschaft (DFG, German Research Foundation) – project numbers 169331223, 388159768. Research in the Wachten lab was supported by grants from the DFG – SFB 1454 – project number 432325352, TRR83 – 112927078, TRR333/1 – 450149205, under Germany’s Excellence Strategy – EXC2151 – 390873048, SPP1926, SPP1726, FOR2743 as well as intramural funding from the University of Bonn. AP and TG are funded by the DFG (Deutsche Forschungsgemeinschaft) 450149205-TRR333/1 (AP: P10; TG: P11).

## Authors contributions

SK, HT, AJ, PL, BDL, ARAAT, NS, IG, NH, ES, RR, SG, LMV, TG, SN, MPJ, NZJ performed the experiments. SK, HT, AJ, PL, IG, AP, TG, MJ, MG, BDL, CAE, MG, RR, PK contributed discussions and performed training. SK, HT, AJ, PL, BDL, NH, PF, ARA, SN, CMS, TJS, FTW, AP, MJ, DW and JWK conceived the experiments. DW, HT and JWK wrote the manuscript.

## Competing financial interests

The authors declare no conflicting financial interest.

## Materials & Correspondence

Requests should be addressed to Jan-Wilhelm Kornfeld or Dagmar Wachten.

## Supplementary information

**Supplementary Figure S1:**
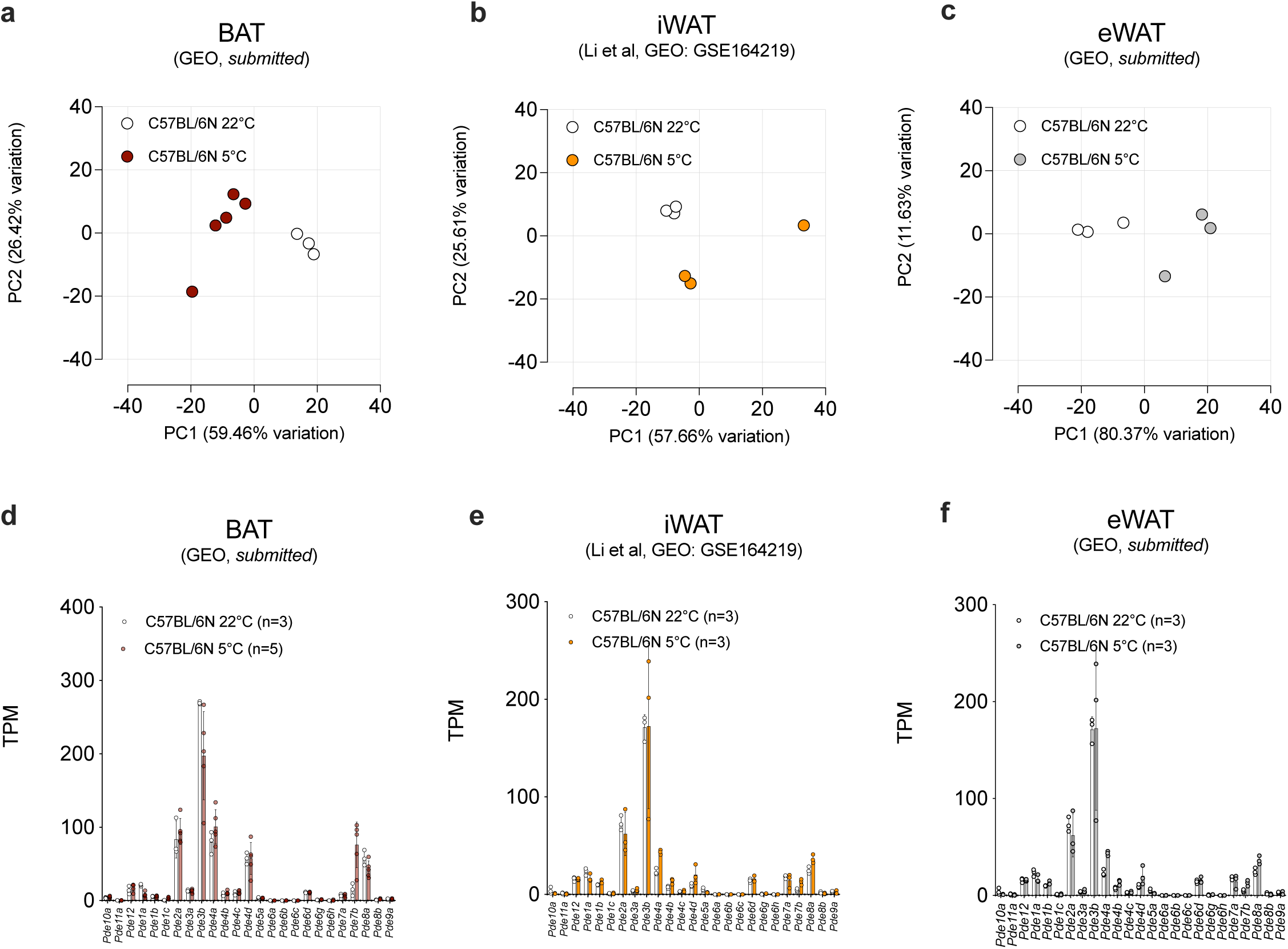
Transcriptional regulation of cAMP biosynthesis in adipose tissue after cold exposure. (**a-c**) Principal component analysis (PCA) of transcriptomic changes in (**a**) BAT, (**b**) iWAT and (**c**) eWAT after 24 h (**a,c**) or 72 h (**b**) of 5° cold exposure. Dots represent individual, chow diet fed male C57BL/6N animals with n=3-5 animals per temperature condition (**d-f**). mRNA expression of phosphodiesterases (PDEs) mRNA abundance in BAT (**d**), iWAT (**e**) and eWAT (**f**) determined by mRNA-seq and depicted in transcripts per million (TPM). (**d-f**) Bar graphs represent mean ± SD with all data points plotted. Unpaired, two-tailed, and non-parametric Mann-Whitney tests were performed to assess statistical significance.

**Supplementary Figure S2:**
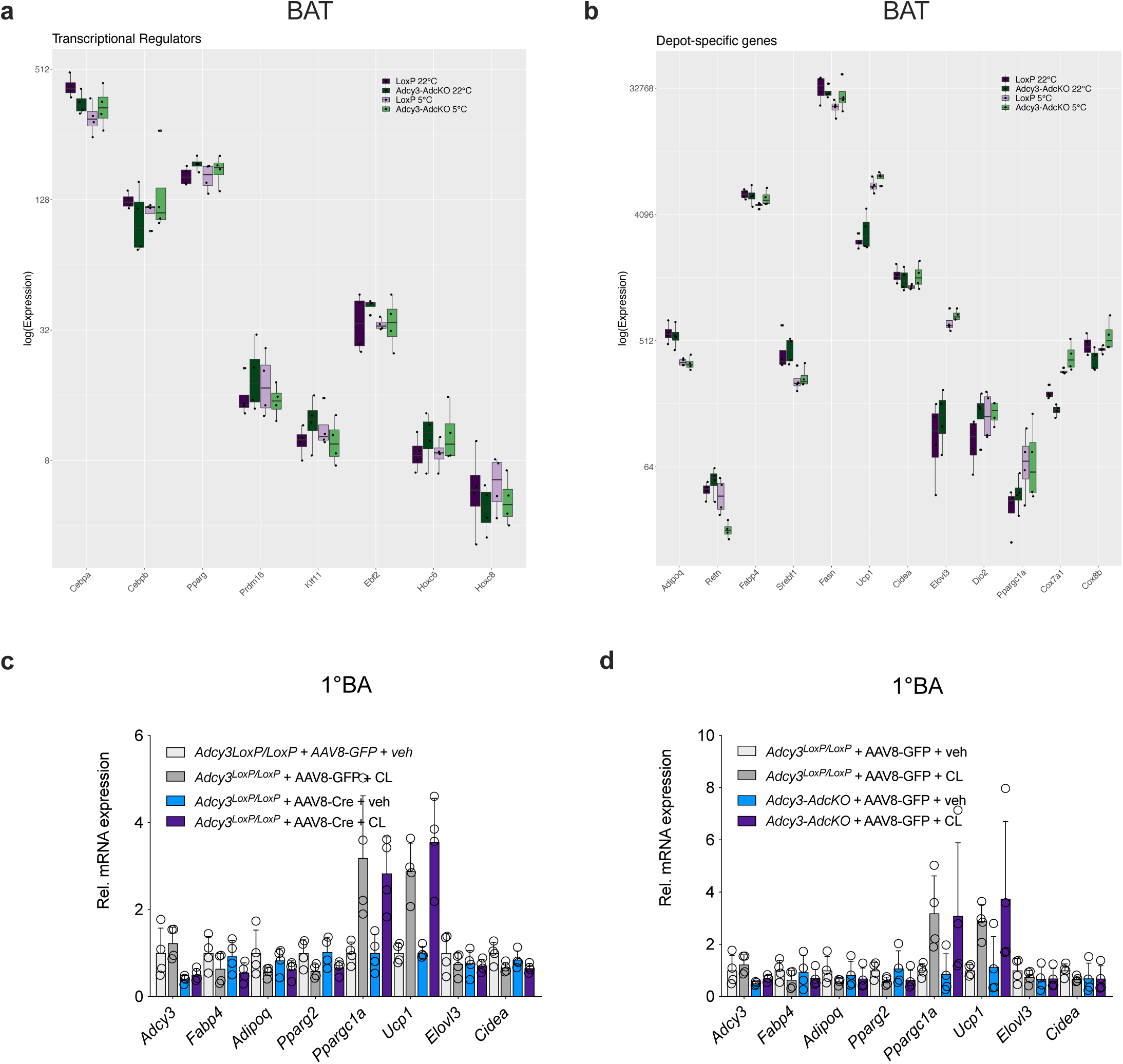
ADCY3 is required for cAMP biosynthesis in BAT and required for cold adaptation in cold. (**a-b**) Expression of indicated (**a**) transcriptional BAT regulator genes and (**b**) adipose depot specific genes in BAT from *LoxP* (n=4 each for 22°C and 5°C) and *Adcy3-AdcKO* mice (n=4 each for 22°C and 5°C) housed at 22°C or after 24 h of 5°C cold exposure as determined by mRNA-seq and depicted in transcripts per million (TPM). (**c-d**) qPCR analysis of indicated BAT mRNA markers measured in 1°BA derived from SVF of *LoxP* animals (**c**) or *LoxP* and *Adcy3-AdcKO* mice (**d**). (**c-d**) Replicates represent n=4 primary brown adipocytes, n=4 mice per adipose depot, stimulation regimen and genotype. Bar graphs represent mean ± SD with all data points plotted. To test for statistical significance, non-parametric (ranked) Kruskal–Wallis one-way analysis of variance (ANOVA) tests with Dunn’s correction for multiple testing were performed.

**Supplementary Figure S3:**
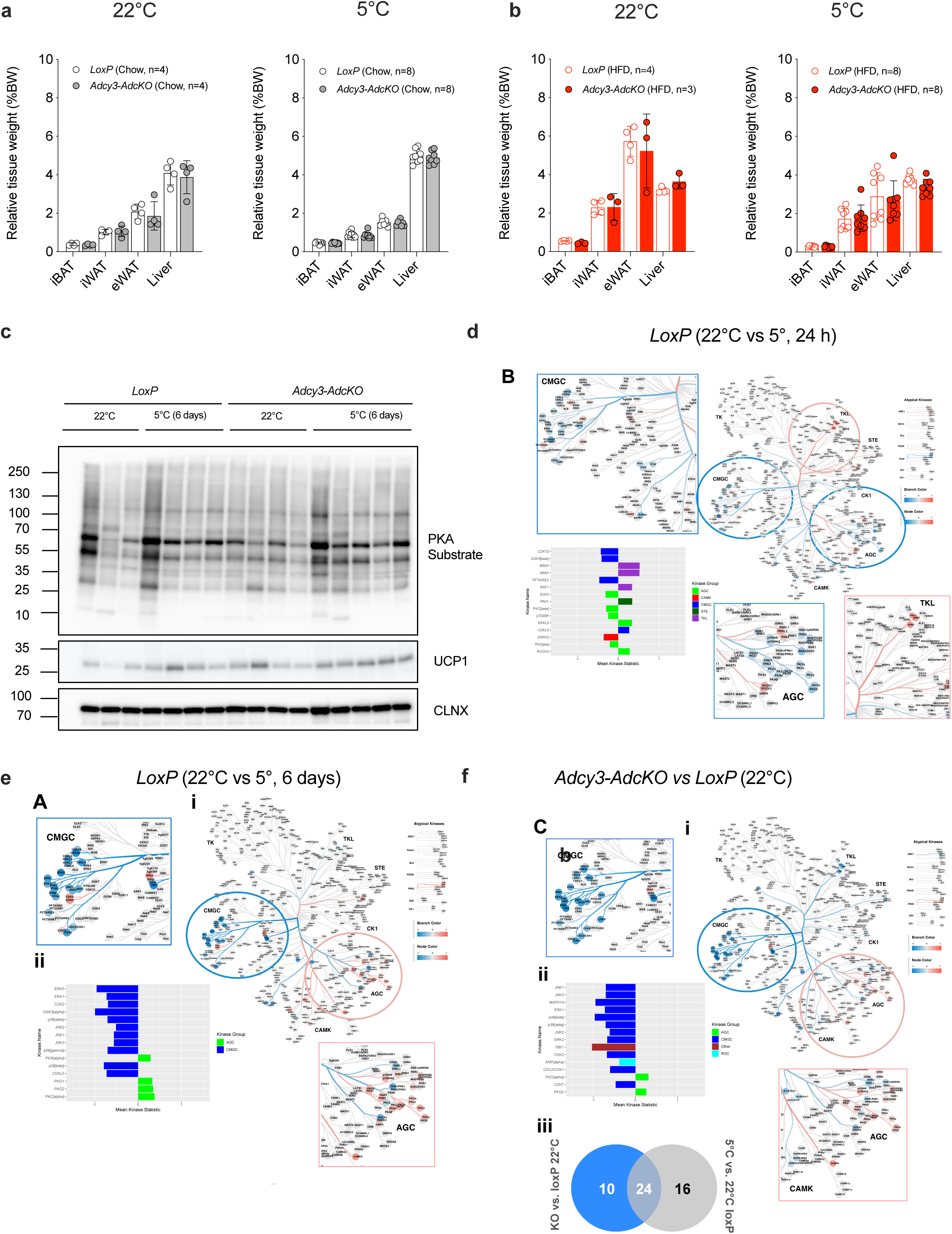
ADCY3 loss in adipocytes predisposes to obesity-associated brown fat dysfunction. (**a-b**) Relative (body-weight corrected) gross tissue weights from indicated adipose tissue depots and liver from *LoxP* (*n=4* for 22°C, n=8 for 5°C) and *Adcy3-AdcKO* mice (*n=4* for 22°C, n=8 for 5°C) fed (**a**) chow diet or (**b**) HFD and housed at 22°C (*left side of panels*) or exposed for 24 h to 5°C (*right side of panels*). Bar graphs represent mean ± SD with all data points plotted. Unpaired, two-tailed, and non-parametric Mann-Whitney tests were performed to assess statistical significance. (**c-d,f**) Analysis of STK activity in murine brown adipose tissue. Kinome trees depict the mean kinase statistic of differentially activated STKs in (**c**) LoxP (22°C vs 5°, 24 h), (**d**) LoxP (22°C vs 5°, 6 days) and (**f**) *Adcy3-AdcKO* vs. *LoxP* (22°C). Inserts show zoom-ins of relevant areas of the kinome tree (blue box: down-regulated kinases; red box: up-regulated kinases). **(c-d,ii**) Mean kinase statistic of 15 most significantly differentially activated kinases is represented in bar graphs. (**f,iii**) Venn diagram depicting differences in differentially activated kinases of the respective comparison. Data represent mean of n=4 biological replicates for all conditions. (**e**) Protein immunoblots of BAT from chow-diet fed, male *LoxP* and *Adcy3-AdcKO* mice housed at room temperature (22°C) or for 6 days in the cold (5°C). Anti-phospho-PKA substrate antibodies were used for detection of PKA phosphorylation substrates. Anti-UCP1 antibodies probed for UCP1 protein levels. anti-HSC70 antibody served as loading control.

**Supplementary Figure S4:**
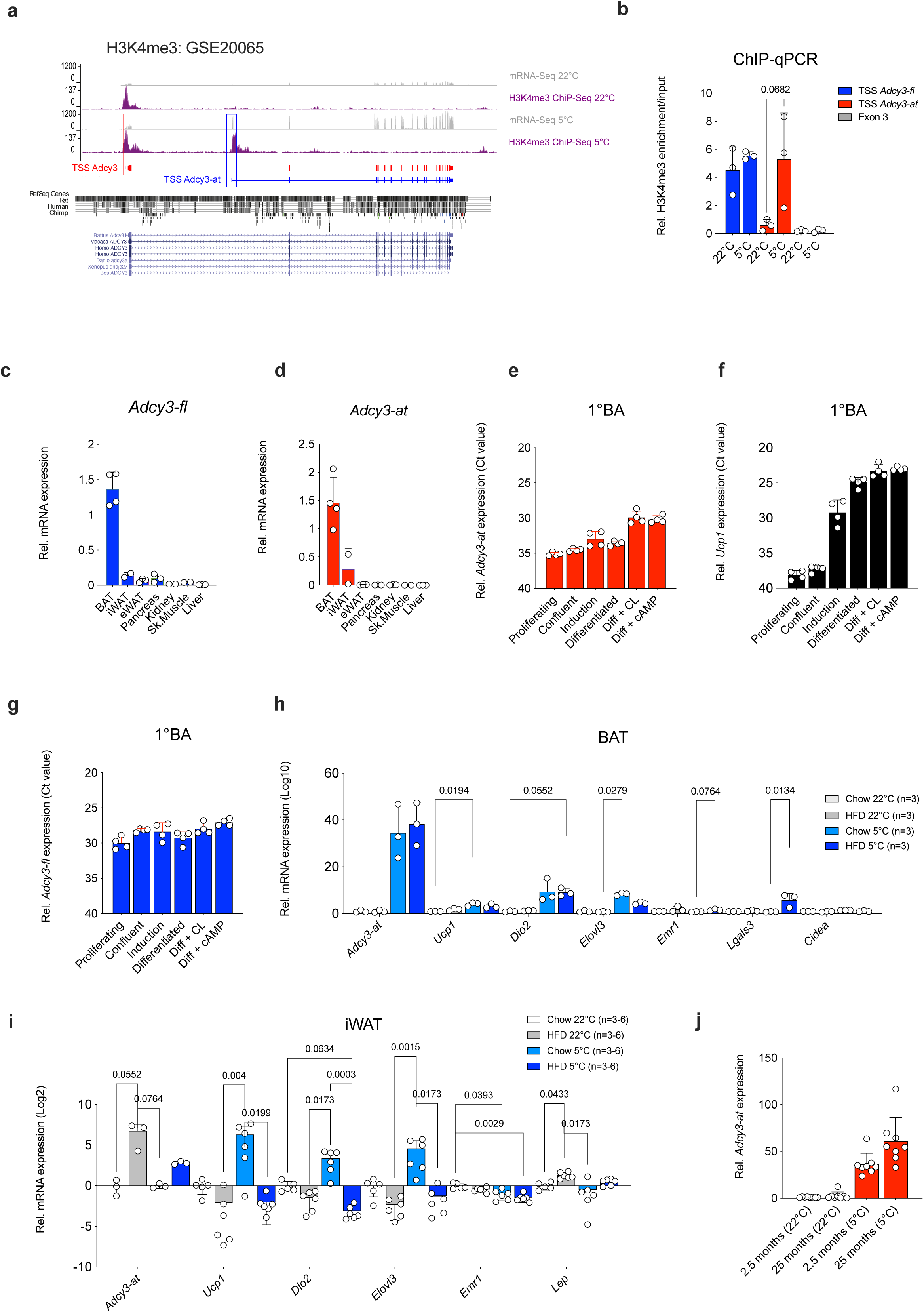
Activated brown adipocytes express a truncated ADCY3 (ADCY3-AT) transcript and protein isoform. (**a**) Genome browser track depicting changes in mRNA expression (grey) and H3K4me3 (magenta) in BAT from chow diet-fed, male C57BL/6N mice housed at 22°C or exposed to 24 h of 5°C cold challenge. Red and blue transcripts depict *Adcy3-fl* and *Adcy3-at* transcripts, respectively, and mRNA isoform specific transcriptional start sites (TSS). (**b**) ChIP-qPCR of H3K4me3 at the TSS of *Adcy3-fl* (blue), *Adcy3-at* (red) and *Adcy3-fl* and *Adcy3-at* common exon 3 (grey). Replicates represent H3K4me3 immunoprecipitations perform in individual animals (n=3). Bar graphs represent mean ± SD with all data points plotted. To test for statistical significance, parametric, unpaired, two-tailed Student’s t-tests were performed. *P-*values are indicated in the panel. (**c-d**) Expression of (**c**) *Adcy3-fl* (blue) and (**d**) *Adcy3-at* (red) in indicated whole tissues from chow diet fed, male C57BL/6N mice. Data points represent individual mice (n=4 BAT, n=2 iWAT, n=3 eWAT, n=3 pancreas, n=3 kidneys, n=2 skeletal muscle, n=3 liver). (**e-g**) Absolute expression (ΔΔCt) of (**e**)*Adcy3-at*, (**f**) *Ucp1* and (**g**) *Adcy3-fl* in BAT-derived SVF at indicated stages of brown adipogenesis and after stimulation with 1 mM db-cAMP or 10 µM CL316,243 as determined by qPCR. Replicates depict experiments performed in SVF isolated from individual mice, i.e., n=4 biological replicates. Bar graphs represent mean ± SD with all data points. (**h**,**i**) Expression of indicated mRNAs in (**h**) BAT or (**i**) iWAT of chow diet or HFD fed male C57BL/6N mice housed at 22°C or after 24 h of 5°C cold exposure as determined by qPCR. Bar graphs represent mean ± SD with all data points plotted. To test for statistical significance, non-parametric (ranked) Kruskal–Wallis one-way analysis of variance (ANOVA) tests with Dunn’s correction for multiple testing were performed. *P*-values are indicated in the panel. (**j**) Expression of *Adcy3-at* in BAT of chow diet fed male C57BL/6N mice at 2.5 months or 25 months of age. Bar graphs represent mean ± SD with all data points plotted.

**Supplementary Figure S5:**
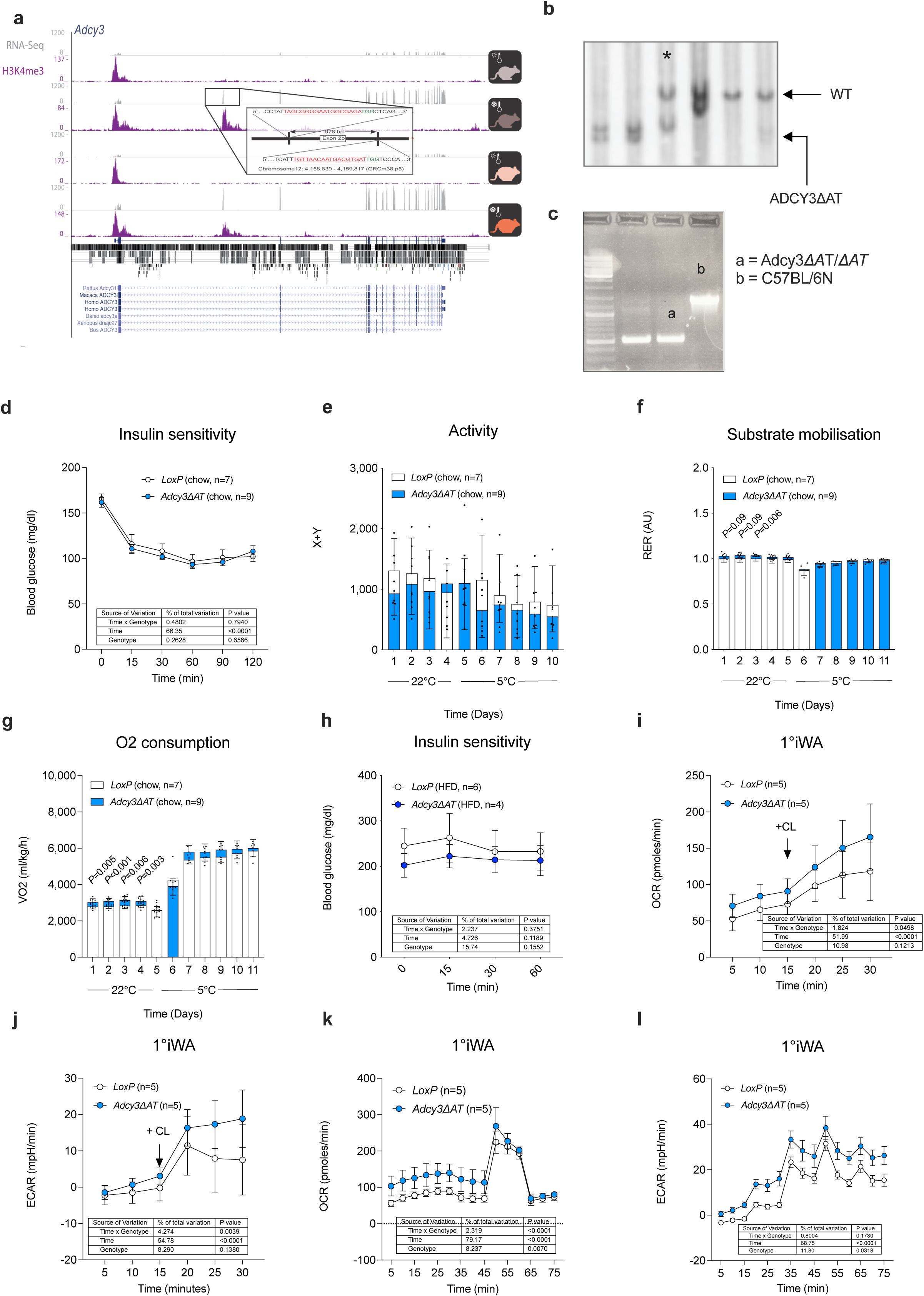
ADCY3-AT inhibits oxidative metabolism *in vitro* and *in vivo*. (**a**) Genome browser track depicting mRNA (grey) and H3K4me3 signals in murine *Adcy3* locus from chow diet fed, male C57BL/6N mice at 22° (light grey) and 5°C (dark grey) as well as HFD-fed, male C57BL/6N mice at 22°C (light orange) and 5°C (dark orange). Location and sequence of sgRNAs used for CRISPR/Cas9 mediated 978-nt deletion *Adcy3-at* TSS are indicated. (**b**) Southern blot analysis of control (C57BL/6N wildtype, WT) and *Adcy3ΔAT* embryonic stem cell DNA. The asterisk depicts a heterozygous *ADCY3ΔAT*/WT ESC. (**c**) PCR from genomic DNA isolated from *Adcy3-at* proficient (WT) and *Adcy3ΔAT* embryonic stem cell DNA. (**d,h**) Blood glucose excursion during insulin tolerance tests performed in (**d**) chow diet and (**h**) HFD fed, male *LoxP* (n=7 chow, n=4 HFD) and *Adcy3ΔAT* (n=9 chow, n=4 HFD) mice. (**e-g**) Quantification of mean (**e**) nocturnal locomotor activity (LMA, **f**) daily respiratory exchange ratios (RER) and (**g**) daily oxygen consumption in chow-diet fed, male *LoxP* (n=7) and *Adcy3ΔAT* (n=9) mice. Bar graphs represent mean ± SD with all data points plotted. Unpaired, two-tailed Student’s t-tests were performed to assess statistical significance. *P*-values are indicated within the panel. (**i-j**) Oxygen consumption rates (**i**) and extracellular acidification rates (**j**) in 1°iWA derived from SVF of *LoxP* or *Adcy3ΔAT* mice after injection with 10 µM CL316243. The numbers of measured wells are indicated in the panel and reflect BA from 2-3 independent mice. (**k-l**) Oxygen consumption rates (**k**) and extracellular acidification rates (**l**) in 1°iWA derived from SVF of *LoxP* or *Adcy3ΔAT* mice. OM, oligomycin; FCCP, AA/Rot, Antimycin A plus Rotenone. The numbers of measured wells are indicated in the panel and reflect BA from 2-3 independent mice. (**m-o**) Bar graphs represent mean ± SD with all data points plotted. Statistical significance was determined by performing Two-Way ANOVA with repeated measurements for x values (mixed models). *Post-hoc P*-value correction to account for multiple testing was performed using Bonferroni adjustment. The source of variation, % of the variation and exact *P* values are given in the table insets. (**d,h-l**) Bar graphs represent mean ± SD with all data points plotted. Statistical significance was determined by performing Two-Way ANOVA with repeated measurements for x values (mixed models). *Post-hoc P*-value correction to account for multiple testing was performed using Bonferroni adjustment. The source of variation, % of the variation and exact *P* values are given in table insets.

**Supplementary Figure S6:**
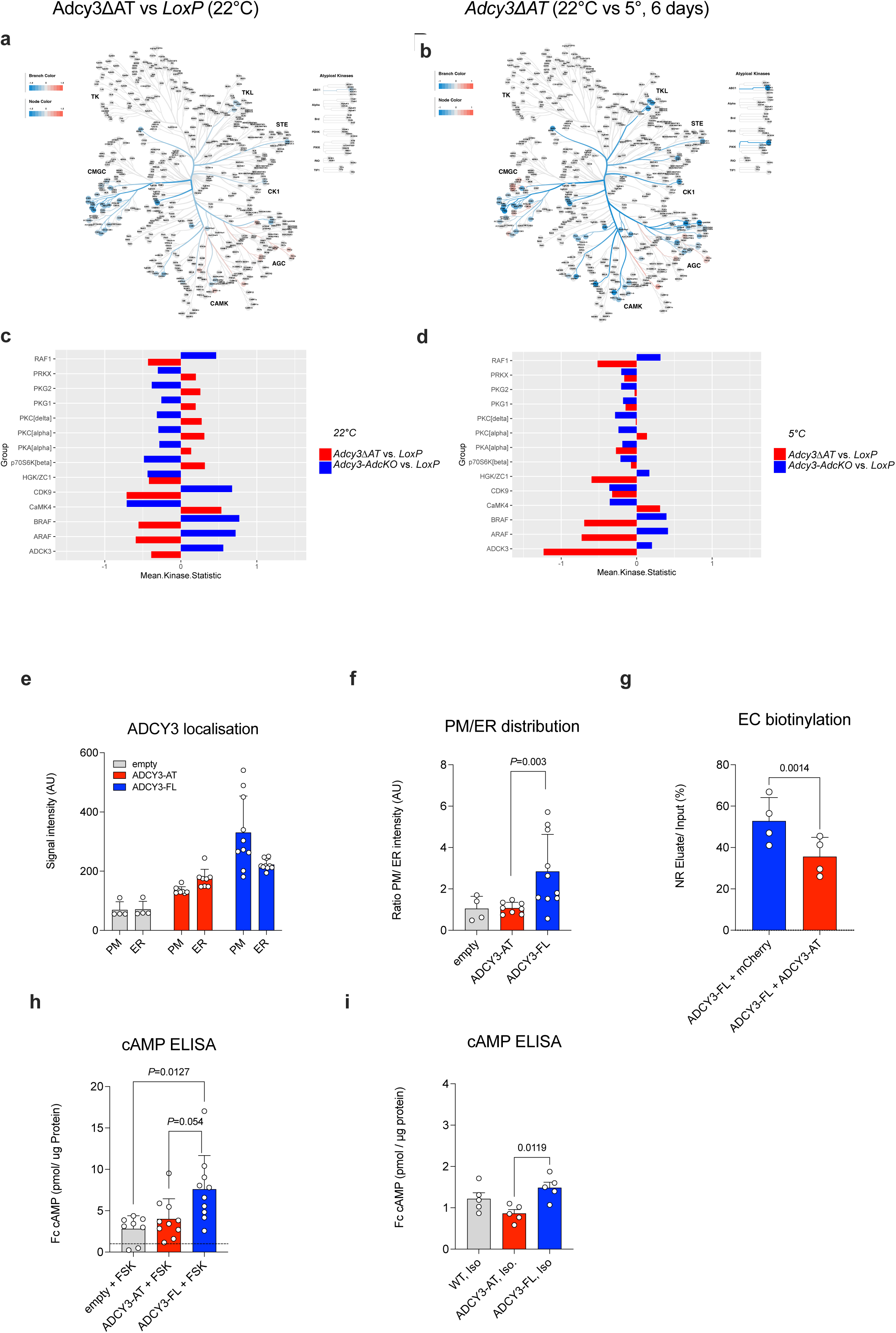
ADCY3-AT alters ADCY3 subcellular localization and limits cAMP biosynthesis. (**a-b**) Analysis of STK activity in *Adcy3ΔAT* BAT. Kinome trees depict the mean kinase statistic of differentially activated STKs in (**a**) *Adcy3ΔAT* vs. *LoxP* (22°C) and (**b**) *Adcy3ΔAT* (22°C) vs. *Adcy3ΔAT* (5°C, 6 days). (**c-d**) Comparison of *Adcy3ΔAT* vs LoxP at 22°C (**c**) and *Adcy3ΔAT* vs *LoxP* after 6 days at 5°C (**d**). At 22°C, a set of kinases is inversely regulated in BAT of *Adcy3ΔAT* and *Adcy3AdcKO* mice compared to *LoxP* mice. The mean kinase statistic of these kinases is shown for 22°C (**c**) and for 5°C (**d**). Data includes n = 4 biological replicates for all conditions. (**e**) Quantification of the average HA-signal intensity within the PM (co-localization with mCherry) and the ER (co-localization with calnexin). Data shown as mean + SD (empty vector: n=4; ADCY3-AT: n=8, ADCY3-FL: n=10). (**f**) Ratio PM/ER distribution (empty vector: n=4; ADCY3-AT: n=8, ADCY3-FL: n=10). Bar graphs represent mean ± SD with all data points plotted. An unpaired, two-tailed, and non-parametric Mann-Whitney tests were performed to assess statistical significance. If relevant, *P*-values are indicated within the panel. (**g**) Quantification of extracellular biotinylation of ADCY3-FL when co-expressed with mCherry or ADCY3-AT (mCherry: n=4, ADCY3-AT: n=4). (**h**) Quantification of intracellular cAMP levels using ELISA. CHO cells stably expressing ADCY3-AT or ADCY3-FL, or empty vector transfected CHO cells were stimulated with DMSO or 2 µM Forskolin for 30 min. Values were normalized to DMSO control and normalized to the respective amount of total protein (empty vector: n=4; ADCY3-AT: n=5, ADCY3-FL: n=5) (**i**) Quantification of intracellular cAMP levels using ELISA. CHO cells stably expressing ADCY3-AT or ADCY3-FL, or empty vector transfected CHO cells were stimulated with buffer or 1 µM Isoproterenol for 15 min. Values were normalized to the respective buffer control and normalized to the respective amount of total protein (empty vector: n=4; ADCY3-AT: n=5, ADCY3-FL: n=5) (**e-i**) Data is shown as mean ± SD with all data points plotted. To test for statistical significance, non-parametric (ranked) Kruskal–Wallis one-way analysis of variance (ANOVA) tests with Dunn’s correction for multiple testing were performed. *P*-values are indicated in the panel.

**Supplementary Figure S7:**
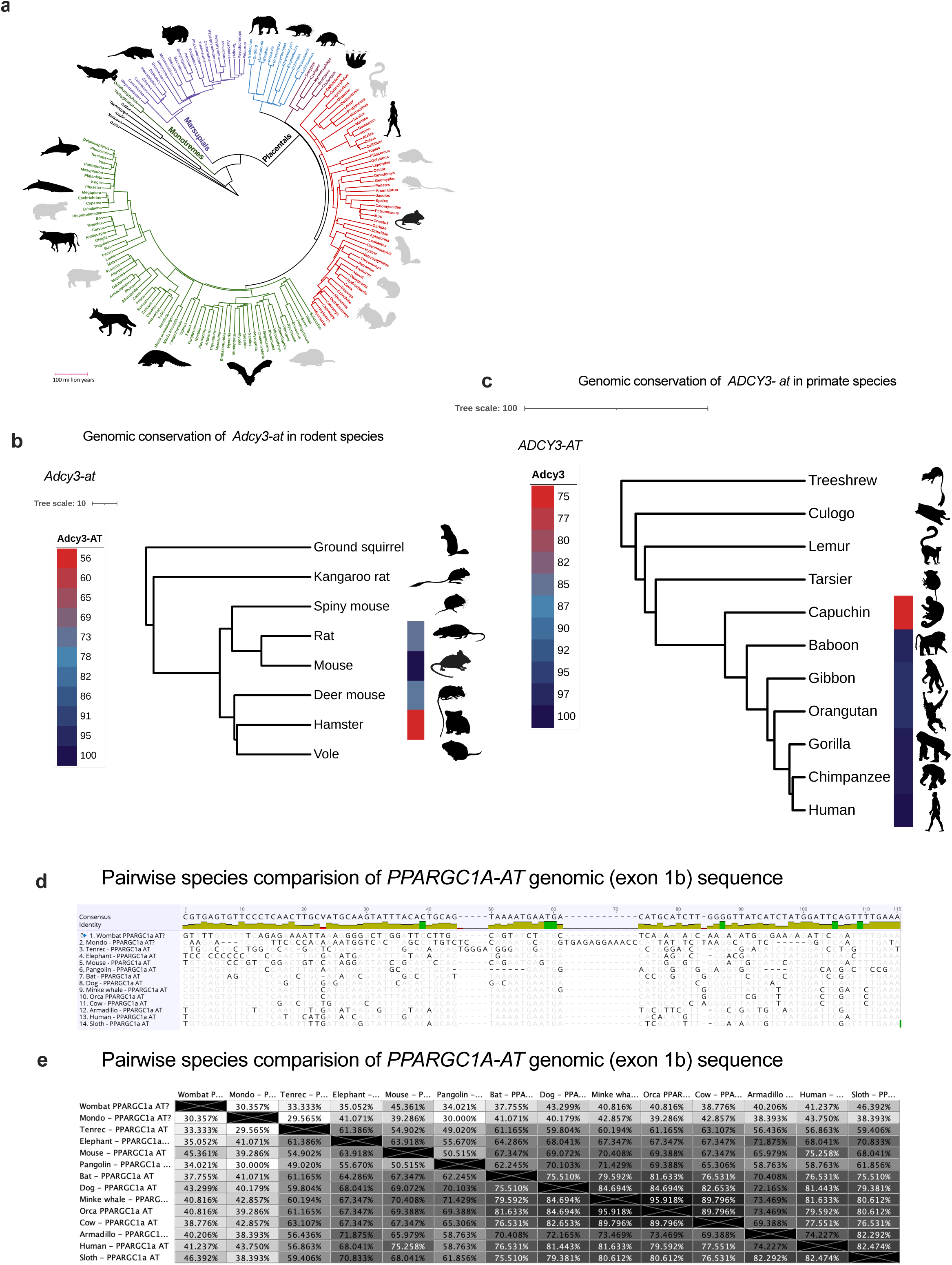
Conserved and cold-inducible PPARGC1A-AT drives *Adcy3-at* expression in BAT. (**a**) Mammalian phylogeny from Meredith et al ^77^ modified in iTOL^89^, with silhouettes in black indicating representative species that were selected for comparative analyses of PPARGC1A-AT. (**b**) Genomic conservation among representative rodents of *Adcy3-at* identified in the mouse. Color gradient indicates percent nucleotide identity of the *Adcy3-at* relative to the mouse. Phylogeny based on Swanson et al ^91^. (**c**) Genomic conservation among representative primates of *Adcy3-at* identified in the human. Color gradient indicates percent nucleotide identity of *Adcy3-at* relative to the human. Phylogeny based on Perelman et al ^90^. (**d**) Nucleotide sequence alignment of *Ppargc1a-at* (exon 1b) from various mammalian species. (**e**) Percent identity matrix of *Ppargc1a-at* among 14 representative mammals (exon 1b).

## Supplementary tables

Supplementary Table S1: Adipose depot-specific RNA-seq from cold-exposed C57BL/6 mice.

Supplementary Table S2: Gene accession numbers for species conservation analyses.

Supplementary Table S3: List of sgRNAs and primers for the generation of *Adcy3*ΔAT mice.

Supplementary Table S4: List of SYBR qPCR primers

Supplementary Table S5: List of locked nucleic acid (LNA) GapmeR inhibitors.

## Notes

### Competing Interest Statement

The authors have declared no competing interest.

### Summary of Updates

Additional method details pertaining to the nature of human brown adipocytes were included.

